# Estrogen-related receptor gamma is required for normal auditory innervation and is essential for hearing

**DOI:** 10.64898/2026.05.02.722410

**Authors:** Shri Vidhya Seshadri, Neil Ingham, Rhianna R. Mackenzie, Adam J. Carlton, Stuart L. Johnson, Darya Alcock, Anwen Bullen, Katie E. Smith, Walter Marcotti, Karen P. Steel, Lisa S. Nolan

## Abstract

Estrogen-related receptor gamma *(ESRRG),* an orphan nuclear receptor with structural homology to the classical estrogen receptors, is widely recognised as a key metabolic regulator involved in mitochondrial, synaptic, and ion-homeostatic pathways. Previous clinical studies suggest a link between ESRRG and auditory function; for example, *ESRRG* has been associated with susceptibility to age-related hearing loss in women and implicated in congenital hearing loss. However, the biological mechanisms by which ESRRG may mediate hearing function remain largely unknown. Here, using a combination of *in vivo* auditory physiological recordings, immunofluorescence analyses, single hair-cell electrophysiology, and transcriptomic approaches, we characterise the phenotype of a new inner-ear conditional *Esrrg* knockout (*Esrrg-*cKO) to investigate the role of *Esrrg* in the auditory system.

We found that *Esrrg-*cKO mice of both sexes develop early-onset hearing loss, as evidenced by elevated auditory brainstem response thresholds and reduced wave 1 amplitudes from two weeks of age. These auditory deficits arise from a combination of early-onset cochlear neuronal and innervation malformations, together with inner hair cell synaptic defects and delayed myelination that persist into adulthood. Furthermore, distortion product otoacoustic emissions and endocochlear potential recordings are normal in *Esrrg*-cKO mice, and although sensory hair cells are preserved, IHCs retain immature biophysical properties. These findings are consistent with auditory neuropathy, and together with our comparative transcriptome analyses, indicate that *Esrrg* is an essential molecular driver of normal cochlear innervation and maturation.

## Introduction

Estrogen-related receptor gamma (*ESRRG; NR3B3*) encodes an orphan nuclear receptor (Hong et al., 1999), a transcription factor closely related to the *classical* estrogen receptors (ER) that, together with highly homologous genes *ESRRA* (*NR3B1*) and *ESRRB* (*NR3B2*), constitutes the NR3B subgroup of the nuclear receptor superfamily (Giguere, 2002). All three estrogen-related receptors (ERRs) are widely recognised as key regulators of energy metabolism, with distinct yet overlapping expression patterns in tissues of high metabolic demand (Tremblay and Giguere, 2007, Huss et al., 2015, Scholtes and Giguere, 2022).

*ESRRG,* the last member of the ERR family to be identified (Hong et al., 1999), is highly expressed across many tissues including the heart, stomach, kidney and brain (Alaynick et al. 2007; Alaynick et al. 2010; Rangwala et al. 2010; Pei et al. 2015; Fox et al. 2022). In mice lacking *Esrrg,* transcriptomic studies have shown that it plays a crucial role in coupling energy metabolism with organ maturation (Alaynick et al., 2007, Dufour et al., 2007, Alaynick et al., 2010, Murray et al., 2013, Yoshihara et al., 2016). In the heart, this role manifests as a metabolic switch that enables the transition from maternal glucose during fetal life to lipid consumption via oxidative metabolism at birth, a process essential for maintaining contractile function (Alaynick et al., 2007, Dufour et al., 2007). Similarly, in skeletal muscle, by maintaining oxidative capacity, *Esrrg* drives myotube differentiation and maturation (Murray et al., 2013). *Esrrg* has also been shown to regulate mitochondrial and synaptic pathways (Pei et al. 2015; Fox et al. 2022; Fox et al. 2025), and to support spatial learning and memory (Pei et al., 2015).

Genetic variants in the *ESRRG* gene have been linked with susceptibility to various metabolic diseases, including osteoporosis (Kim et al., 2025), diabetes (Rampersaud et al., 2007, Kim et al., 2017), and obesity (Dong et al., 2016, Daily et al., 2019). Furthermore, in the auditory system, *ESRRG* has been associated with susceptibility to age-related hearing loss (ARHL) in women of post-menopausal age (Nolan et al., 2013), and a chromosomal deletion disrupting *ESRRG* has been implicated in a monogenic form of congenital hearing loss (Schilit et al., 2016). However, despite the connections between *ESRRG* and auditory function, the molecular mechanisms by which *ESRRG* mediates hearing remain poorly understood.

*Esrrg* is expressed early in the developing auditory system (Hermans-Borgmeyer et al., 2000, Patthey et al., 2016). *In-situ* hybridization studies in mice have shown expression is detectable at embryonic day (E)10.5 in the otic vesicle, from which the inner ear structures arise, with robust expression evident by E16.5 in both vestibular and cochlear ganglia (Hermans-Borgmeyer et al., 2000). Immunolabelling studies have also reported ESRRG expression in cochlear spiral ganglion neurons (SGN), as well as in discrete supporting cell types and inner hair cells (IHC) (Nolan et al., 2013). Consistent with this, recent bulk and single-cell RNA-sequencing studies have detected *Esrrg* in both developing (Li et al., 2020) and adult murine SGNs (Petitpre et al., 2018).

The SGNs are central for propagating acoustic information to the brain. In humans and other mammals, there are two main types. Type I SGNs constitute the vast majority (90-95%) of the afferent innervation and receive input from IHCs, the primary mechanotransducers of the cochlea (Coate et al., 2019, Johnson et al., 2019). Together, Type I SGNs and the IHCs they innervate are fundamental for reception of sound and encoding the full dynamic range of hearing. Type II SGNs, which comprise the remaining afferent innervation (5-10%), innervate outer hair cells (OHCs), and are thought to function as cochlear nociceptors (Flores et al., 2015, Liu et al., 2015).

Previously, we showed that mice carrying a germline knockout (KO) of the *Esrrg* gene develop hearing loss with female mice exhibiting worse hearing than males (Nolan et al., 2013). However, most *Esrrg* KO mice die during the early postnatal period, with very few surviving into adulthood (Alaynick et al., 2007, Alaynick et al., 2010, Nolan et al., 2013), limiting mechanistic insight into the role of *Esrrg* in the auditory system. Here, we report the phenotypic investigation of a new conditional knockout mouse model, *Esrrg-*cKO, in which *Esrrg* is specifically disrupted in the inner ear. We show that *Esrrg*-cKO mice exhibit reduced numbers of Type I SGNs at birth, and that the bundling of peripheral radial fibres is markedly sparse. By the onset of hearing, which in mice occurs at around postnatal day 12 (Mikaelian and Ruben, 1965), *Esrrg*-cKO mice exhibit disrupted IHC ribbon synapses, delayed myelination, and significant hearing impairment. These anomalies persist into adulthood, and although we detected no loss of hair cells, we found that IHCs fail to mature into fully functional sensory receptors. Together with comparative transcriptome analyses and normal distortion product otoacoustic emissions (DPOAE) and endocochlear potentials (EP), our findings support a phenotype consistent with auditory neuropathy and indicate that the molecular signature of *Esrrg* is essential for normal auditory innervation and cochlear maturation.

## Results

### Generation of Esrrg-cKO mice

To characterise the role of *Esrrg* in hearing, we generated a new mouse model using *Sox10-*Cre to disrupt *Esrrg* early in inner ear development (Fig.1A) (Watanabe et al., 2000, Matsuoka et al., 2005). Ensembl reports four protein coding transcripts for the *Esrrg* gene in mice (Fig.1C) (Dyer et al., 2025). *Esrrg*-202 and *Esrrg*-203 encode the same short 48.6kDa isoform whereas *Esrrg*-201 encodes the larger canonical isoform (51kDa) containing a longer N-terminal domain. *Esrrg*-204 is predicted to encode an extra-short isoform of 5.3kDa. qPCR analysis of P1 cochlear tissue with a Taqman probe spanning the floxed exon, which is common to all coding isoforms of the *Esrrg* gene, confirmed significant knockdown of *Esrrg* mRNA in *Esrrg-*cKO mice compared to *Esrrg^fl/fl^* littermate controls (Fig. 1B). Similar to previous reports using this Cre driver (Kochaj et al., 2022, Nolan et al., 2022), a small amount of residual expression of the target gene was detected (Fig. 1B).

**Figure 1.**
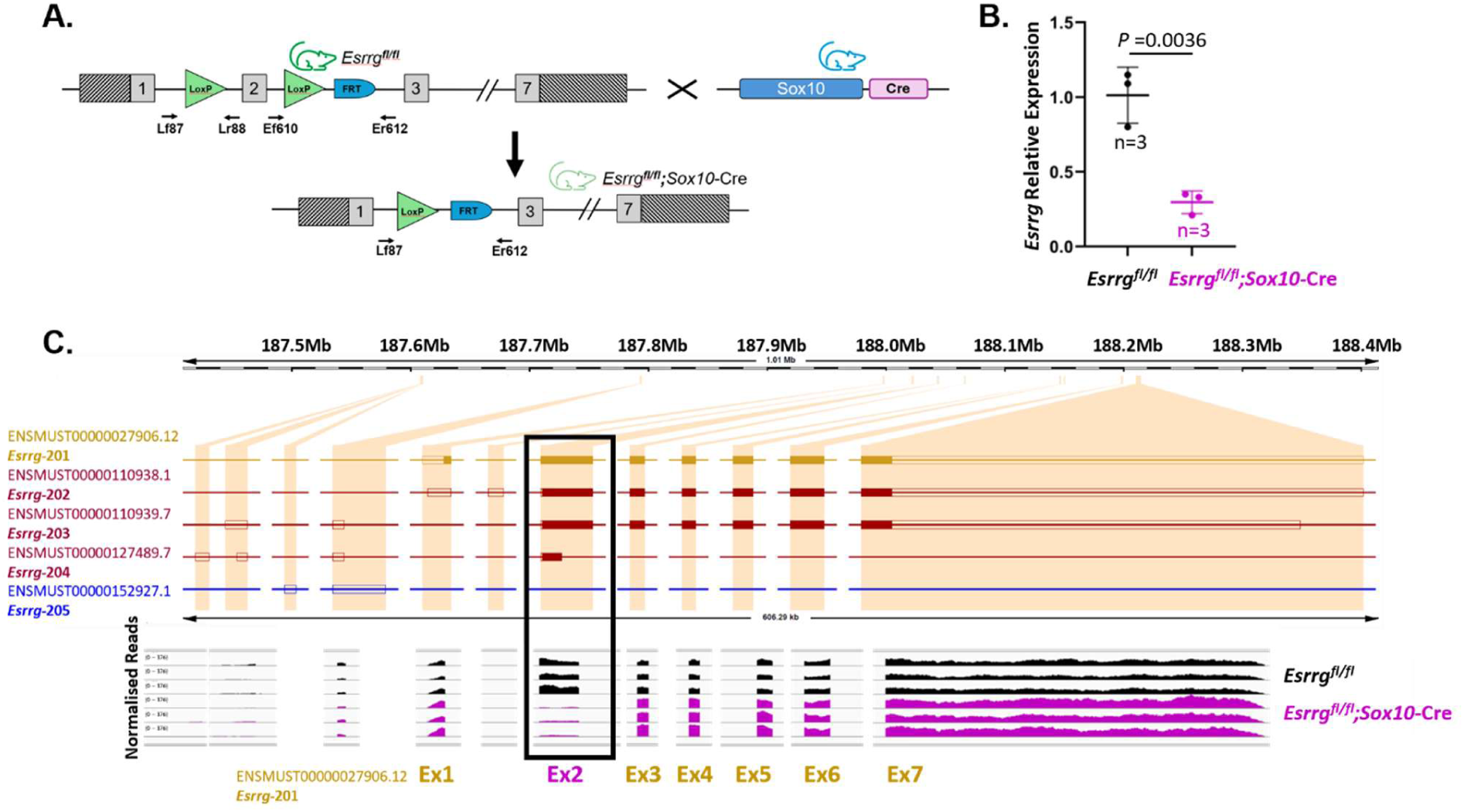
Generation of *Esrrg*-cKO mice. **(A)** Schematic of the mouse canonical sequence for *Esrrg* illustrating the design of the floxed allele and the recombined allele generated in the presence of the *Sox10*-Cre transgene. Coding exons - grey; UTR - diagonal rectangles; LoxP sites (green) flank the critical exon; black arrows - genotyping primers. **(B)** qPCR analysis of *Esrrg* expression in P1 cochlear tissue. Relative quantification levels are adjusted to *Gapdh* as an endogenous control and normalised to *Esrrg*^fl/fl^ levels. Mean (±SD) from female mice is shown; statistical comparisons: unpaired *t*-test. **(C)** Schematic of the mouse genomic locus for *Esrrg* (ENSMUSG00000026610; MGI:1347056; adapted from Ensembl, GRCm38, release 102) to show all four coding transcripts aligned with the normalized reads detected from the bulk RNA-Seq of P1 cochleae tissue. Upper panel: The canonical transcript (*Esrrg*-201) encodes a 51kDa isoform, has 7 exons (shaded rectangles) and part of exons 1 and 7 are untranslated (open rectangles). Transcripts *Esrrg*-202 and *Esrrg*-203 lack the coding sequence of exon 1 and encode a shorter 48.6kDa isoform; transcript *Esrrg-204* only encodes part of exon 2, 5.3kDa. Lower panel: Normalised reads from 3 representative *Esrrg^fl/fl^;Sox10-*Cre mice (magenta) and 3 *Esrrg*^fl/fl^ littermate controls (black); the boxed region shows the critical exon common to all coding transcripts is significantly reduced in *Esrrg^fl/fl^;Sox10-*Cre mice. Data shown is from female mice.

We then examined the RNA-Seq reads captured at the *Esrrg* locus as part of our transcriptome analysis of whole cochleae tissue in *Esrrg-*cKO mice at P1 (Fig.1C, Fig.7). Consistent with our qPCR analysis, this confirmed significant knockdown of the floxed exon (Fig.1C). No reads mapped to the locus analogous to the 2^nd^ untranslated exon of transcript *Esrrg*-202 nor the 1^st^ untranslated exon of transcript *Esrrg*-204 suggesting these transcripts are not present in the cochlea. In comparison, reads were detected for the 5’ untranslated exons of *Esrrg*-203, and for the subsequent exons downstream of the critical exon (Fig.1C). Therefore, we deduced that both the canonical transcript, *Esrrg*-201, and transcript *Esrrg*-203 are present in the cochlea, and this conditional knockout strategy selectively disrupts transcription of the critical exon; thus, generating a novel cKO line for *Esrrg* in the inner ear.

### Esrrg-cKO mice exhibit early-onset hearing loss

Auditory brainstem response (ABR) recordings capture the synchronous response of auditory neurons to sound (Ingham, 2019). To assess auditory function in *Esrrg*-cKO mice, we performed ABRs at key timepoints: 2 weeks [shortly after hearing onset (Mikaelian and Ruben, 1965)], 4 weeks [when auditory afferent innervation is mature (Coate et al., 2019)], and 9 weeks of age [young adulthood, when female mice are sexually mature (Pritchett and Taft, 2007)] (Fig. 2A-C). In control mice (*Esrrg^+/+^*), ABR thresholds from click stimuli and 3-12kHz pure tones improved from 2 to 4 weeks of age (Fig. 2A-B; S1 Table), consistent with the ongoing development of auditory innervation (Coate et al., 2019). Thresholds then remained stable between 4 and 9 weeks (Fig. 2B,C; S1 Table). In comparison, in *Esrrg*-cKO mice, pure-tone evoked ABR thresholds were significantly elevated across 6-36 kHz from 2 weeks (Fig. 2A,B; S1 Table). By 4 weeks, thresholds to click stimuli were significantly raised and the pure tone responses, although showing some improvement across most frequencies, remained significantly elevated in the low-mid frequencies (3-24kHz; Fig. 2A,B; S1 Table). Little additional deterioration in thresholds was evident across any stimuli by 9 weeks (Fig. 2C; S1 Table). We then stratified our results by sex to assess for potential sex-specific effects we but did not find any effect of sex on the raised ABR thresholds (Fig.S1; Table S2). These results suggest that *Esrrg* is important for the normal development of auditory function in both males and females.

**Figure 2.**
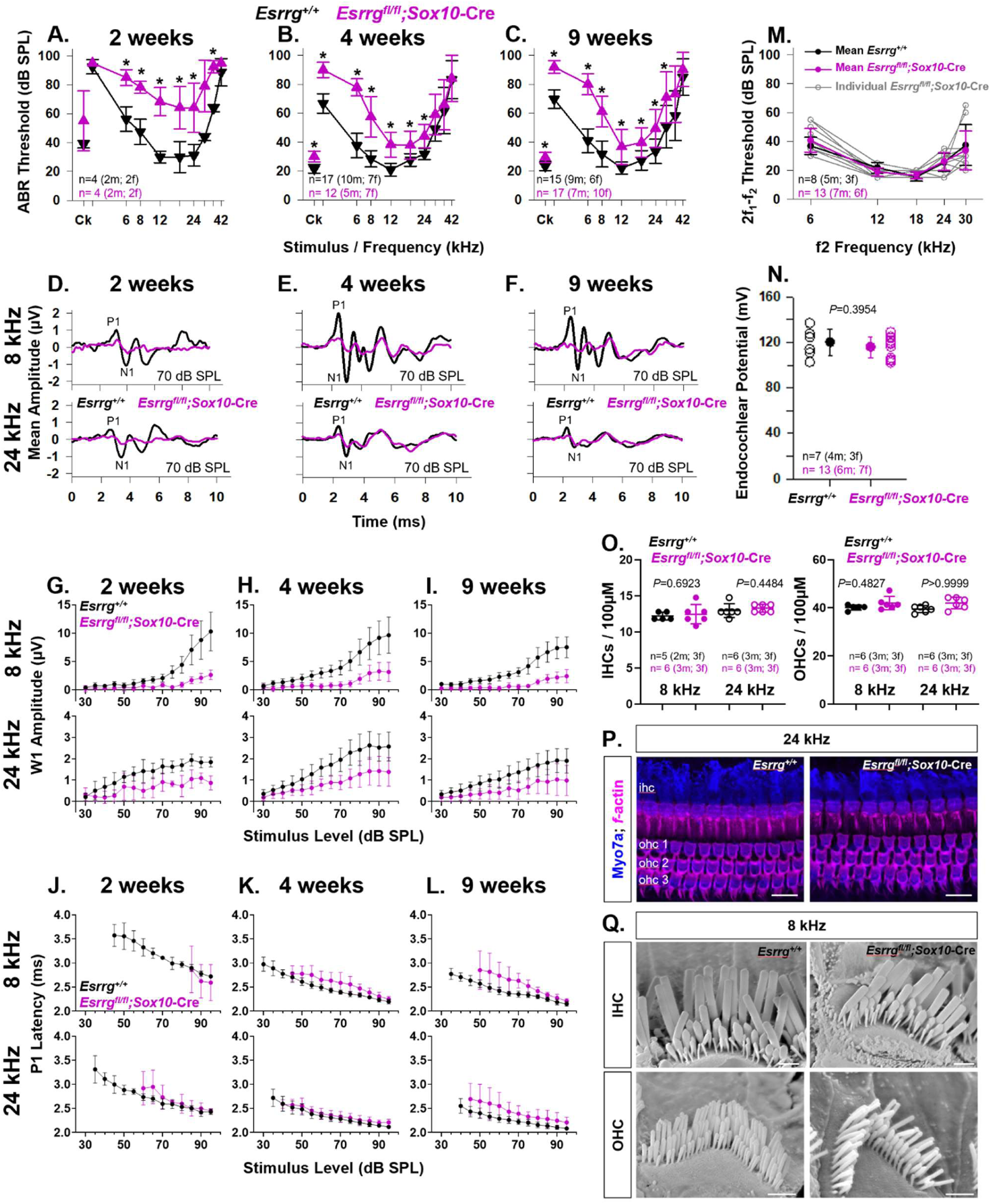
ABR thresholds are elevated in *Esrrg*-cKO mice in the absence of sensory hair cell loss. **(A-C)** Mean ABR thresholds (±SD) to click and pure tone stimuli in *Esrrg^+/+^* and *Esrrg^fl/fl^;Sox10-*Cre mice at 2 (P14), 4 (P31-P37) and 9 (P60-P64) weeks of age. The number and sex of the mice is shown below the thresholds. Statistical comparisons: unpaired *t-*test with Holm-Šídák’s correction for multiple testing; *denotes significance; for *p(adj)* see Table **S1**. **(D-F)** Mean tone evoked ABR waveforms recorded at 70dB sound pressure level (dB SPL) for 8 kHz and 24 kHz pure tones in *Esrrg^+/+^* and *Esrrg^fl/fl^;Sox10-*Cre mice at 2, 4 and 9 weeks of age. P1 and N1 denote the peak and trough of Wave 1, respectively. **(G-I)** Mean wave 1 amplitude (±SD) at 8 kHz and 24 kHz plotted as a function of dB sound pressure level in *Esrrg^+/+^* and *Esrrg^fl/fl^;Sox10-*Cre mice at 2, 4 and 9 weeks of age. For clarity, data is shown from 30dB SPL. **(J-L)** Mean wave 1 latency (±SD) at 8kHz and 24kHz plotted as a function of dB sound pressure level in *Esrrg^+/+^* and *Esrrg^fl/fl^;Sox10-*Cre mice at 2, 4 and 9 weeks of age. Latency is shown from the mean threshold (dB SPL) for each genotype and frequency. Data from (D-L) were obtained from the same mice as in (A-C). **(M)** Distortion product otoacoustic emissions (DPOAE) in *Esrrg^+/+^* and *Esrrg^fl/fl^;Sox10-*Cre mice at 9 weeks of age (P57-P62). Mean (±SD) 2f_1_-f_2_ thresholds plotted as a function of f2 frequency (kHz) do not significantly differ between the two genotypes (Statistical comparisons: *p* = 0.8621, 2-way ANOVA with Tukey’s multiple correction. **(N)** Endocochlear potential (EP) recordings from *Esrrg^+/+^* and *Esrrg^fl/fl^;Sox10-*Cre mice at 9 weeks of age (Statistical comparisons: unpaired *t-*test). Recordings were taken from the same mice used in (M); number and sex are shown below the thresholds. (M,N) Solid circles: mean value (±SD); open circles: single recordings. **(O)** Mean number (±SD) of IHC and OHC per 100µm length of the organ of Corti at 8 kHz and 24 kHz cochlear regions in *Esrrg^+/+^*and *Esrrg^fl/fl^;Sox10-*Cre mice at 9 weeks of age (Statistical comparisons: unpaired *t*-test). **(P)** Representative maximum intensity projections of confocal z-stacks of cochlear whole mount preparations from *Esrrg^+/+^* and *Esrrg^fl/fl^;Sox10-*Cre mice at 9 weeks of age immunolabelled with the hair cell marker Myosin7a (Blue), co-stained with Phalloidin to *f-*actin (Magenta) to reveal the hair bundles and cuticular plate Pillar cell surfaces also label. Scale bar: 20µm. Counts were taken from the same mice used in (C). **(Q)** Representative scanning electron micrographs of the apical inner and outer stereociliary hair bundles in *Esrrg^+/+^* and *Esrrg^fl/fl^;Sox10-*Cre mice at 9 weeks of age. Scale bar: 1µm.

#### Loss of characteristic ABR waveform shape

The ABR is defined by a characteristic waveform of which there are typically 4-5 detectable peaks in mice (Willott, 2006, Ingham, 2019). Defects in ABR wave 1 can sometimes indicate a defect at the IHC ribbon synapse (synaptopathy) and/or the peripheral auditory myelination (Liberman, 2017, Wan and Corfas, 2017). We compared the mean tone-evoked ABR waveforms for 8 kHz (mid-apical) and 24 kHz (mid-basal), frequencies that showed substantial and small threshold shifts from 4 weeks, respectively (Fig. 2B,D-F). Consistent with the ongoing maturation of auditory thresholds (Fig. 2A,B), control mice displayed a mature ABR waveform by 4 weeks at both 8 kHz and 24 kHz (Fig. 2D,E). Conversely, in *Esrrg*-cKO mice, the entire ABR waveform was severely disrupted across both frequencies from 2 weeks, with only some clearer detection of wave 4 apparent by 4 weeks (Fig. 2D-E). Plotting wave 1 amplitude and latency as a function of the sound pressure level confirmed a substantial reduction in wave 1 amplitude from 2 weeks across both frequencies (Fig. 2G-I) in conjunction with a small increase in latency evident at both frequencies from 9 weeks (Fig. 2J-L). No effect of sex was found in *Esrrg*-cKO mice when these results were stratified by sex (Fig. S2). Wave 1 amplitude was greatly reduced in both sexes at each time point with the clearest elevation in latency evident in both sexes from 9 weeks (Fig. S2).

#### Normal DPOAEs, EPs and auditory ossicles

Next, we recorded DPOAEs, a natural consequence of OHC somatic electromotility and thus, a measure of OHC function (Kemp, 2002). However, DPOAE thresholds (2f_1_-f_2_) in young adult (9 weeks old) *Esrrg*-cKO mice did not significantly differ from control mice across any f2 frequency (6-30kHz) assessed (Fig. 2M; *p* = 0.8621, 2-way ANOVA). Similarly, 2f_1_-f_2_ amplitudes plotted as a function of f2 sound pressure level were also indistinguishable from control mice across all frequencies (Fig. S3A) suggesting normal OHC function in *Esrrg*-cKO mice. *Esrrg* is known to have a role in potassium ion homeostasis in many tissues (Alaynick et al., 2010, Luo et al., 2013, Wang et al., 2024). Thus, we then investigated whether the EP, the driving force underlying mechanosensory transduction and measure of the metabolic activity of the stria vascularis was reduced in *Esrrg*-cKO mice, but we found no significant difference in EP recordings between genotypes (Fig. 2N; 120.2±11.53 mV versus 116.0 ±9.29 mV *p*=0.3954, unpaired *t*-test, in *Esrrg ^+/+^* versus *Esrrg*-cKO mice, respectively). Given the elevated ABR thresholds (Fig. 2A-C), particularly in the low frequencies, we also investigated whether the morphological characteristics of the auditory ossicles were normal but found no difference between genotypes nor sex, suggesting normal middle ear function (Fig. S4). Similarly, examining DPOAE and EP recordings by sex did not uncover any sex differences in *Esrrg*-cKO mice(Fig. S3B-D).

#### No loss of sensory hair cells

To better understand the nature of the hearing loss in *Esrrg*-cKO mice we turned our attention to the sensory hair cells focusing on 8 kHz and 24 kHz in line with our waveform analysis. Quantifying the number of IHCs and OHCs at 9 weeks old with the hair cell marker Myosin 7a in conjunction with the fluorescent hair bundle dye Phalloidin to label *f-*actin did not reveal any loss of sensory hair cells and the structural integrity of the organ of Corti appeared normal (Fig. 2O,P). Scanning electron microscopy (SEM) also confirmed the presence of the normal staircase arrangement of the stereociliary hair bundles (Fig. 2Q). Collectively, the absence of sensory hair cell loss in the presence of raised ABR thresholds in conjunction with normal DPOAE and EP recordings suggest that the origin of the hearing loss in *Esrrg*-cKO mice is auditory neuropathy (Shearer and Hansen, 2019).

### Loss of IHC ribbon synapses in Esrrg-cKO mice from postnatal development

To determine if the reduction in ABR wave 1 in *Esrrg*-cKO mice (Fig.2D-I) might be due to a defect at the IHC ribbon synapse, we immunolabelled wholemount preparations of the organ of Corti from adult (Fig. 3A,B) and postnatal (Fig. 3H-K) mice with an antibody directed against C-Terminal binding protein 2 (CtBP2) to label pre-synaptic ribbons (Uthaiah and Hudspeth, 2010) and Glutamate ionotropic receptor AMPA type subunit 2 (GluA2), a marker for the post-synaptic density (Matsubara et al., 1996). Consistent with our waveform analysis (Fig. 2D-I), both mid-apical (8 kHz) and mid-basal (24 kHz) regions of the cochlea were examined. In adult *Esrrg^+/+^* mice, the typical juxtaposition of a pre-synaptic ribbon paired with an adjacent GluA2 post-synaptic density was observed confirming the structural integrity of IHC ribbon synapses (Fig. 3A,A’’’). In comparison, in *Esrrg*-cKO mice many ribbons and GluA2 post-synaptic densities were orphans exhibiting little colocalisation (Fig. 3B,B’’’,C, D) and the number of paired synapses was significantly reduced (Fig. 3B,B’’’,E). The total numbers of ribbons and post-synaptic densities were also significantly reduced (Fig. 3F,G). Examining these results by sex revealed comparable results in *Esrrg*-cKO mice from both males and females (Fig. S5). However, there was a trend for male *Esrrg*-cKO mice to exhibit a greater proportion of orphan ribbons, and this reached significance in the basal cochlear coil (Fig. S5A).

**Figure 3.**
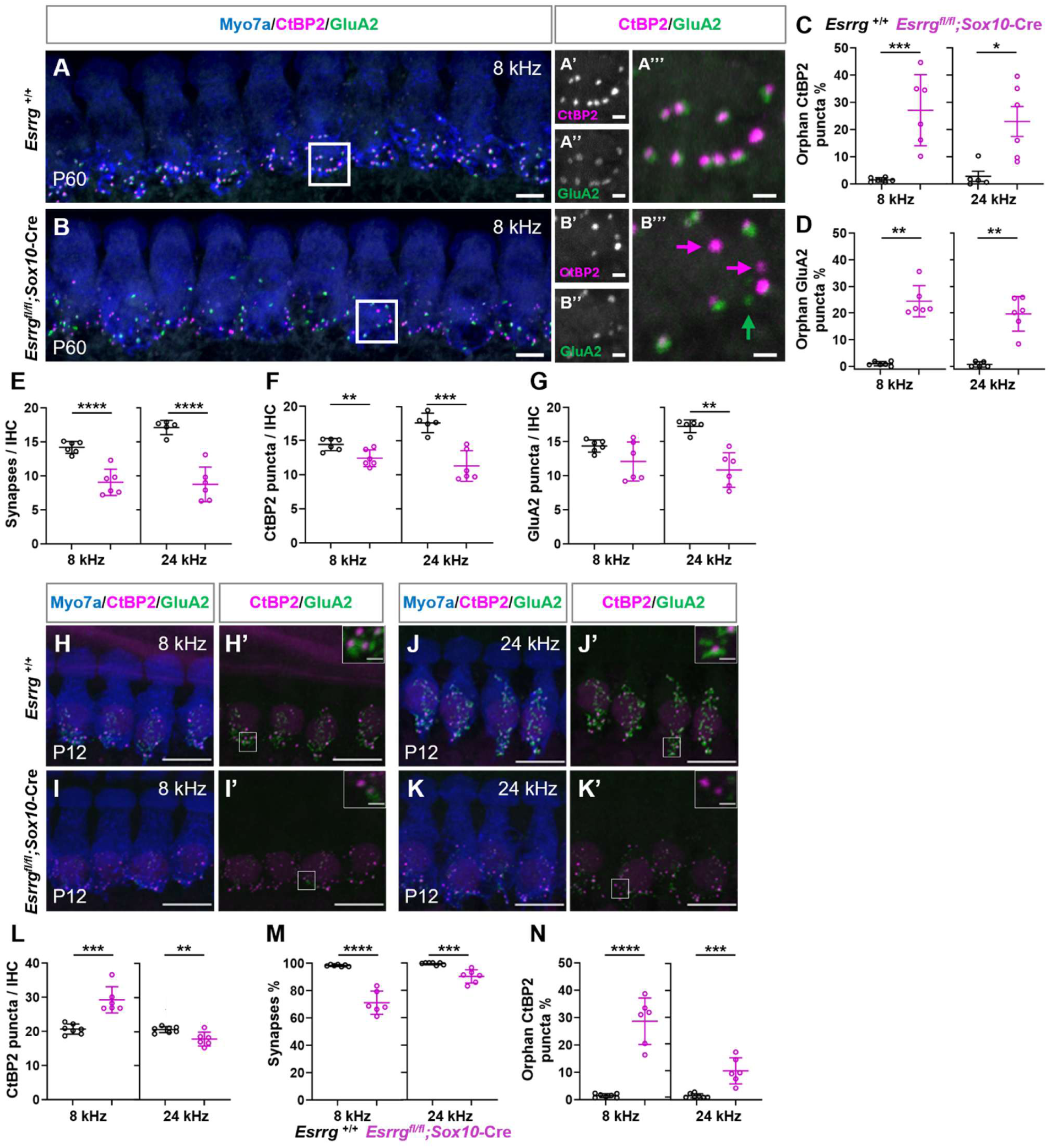
The architecture of the IHC ribbon synapse is disrupted in *Esrrg*-cKO mice. Maximum intensity projections of confocal z-stacks of cochlear whole mount preparations from *Esrrg^+/+^* **(A,H,J)** and *Esrrg^fl/fl^;Sox10-*Cre **(B,I,K)** mice at P60-P64 **(A,B)** and P12 **(H-K)** immunolabelled with antibodies to Myosin7a, CtBP2 and GluA2. In adult *Esrrg^+/+^* mice, pre- (CtBP2) and post- (GluA2) synaptic components colocalise to form paired synapses (zoomed-in insets **A’-A’’’** from white boxed region **A**). In comparison, in adult *Esrrg^fl/fl^;Sox10-*Cre mice many pre- and post-synaptic components are uncoupled (**B’-B’’’**; arrows). **(C)** Orphan ribbons and **(D)** orphan GluA2 densities in adult *Esrrg^+/+^* and *Esrrg^fl/fl^;Sox10-*Cre mice expressed as a percentage of the total number of ribbons / GluA2 densities, respectively. **(E)** Number of paired synapses, **(F)** CtBP2 puncta and **(G)** GluA2 puncta per IHC in adult *Esrrg^+/+^* and *Esrrg^fl/fl^;Sox10-*Cre mice - counts were taken from the same mice used in ABR recordings, Fig. 2C. **(H’-K’)** Contain zoomed-in insets from the adjacent white boxed regions to show weaker GluA2 signal intensity in P12 *Esrrg^fl/fl^;Sox10-*Cre mice compared to controls. Number of CtBP2 puncta **(L)**, paired synapses **(M)**, and orphan ribbons **(N)** per IHC at P12. Paired synapses **(M)** and orphan ribbons **(N)** are expressed as a percentage relative to the total number of ribbons. Data are shown for both the mid-apical (8kHz) and mid-basal (24kHz) cochlear regions. Representative images are from female mice. Data in **(C-G)** and **(L-N)** is sampled from 8-10 and 10-12 IHCs, respectively in at least 2-3 mice per genotype per sex. All data is plotted as mean values ± SD; *p<0.05; **p<0.01; ***p<0.001; ****p<0.0001 unpaired *t*-test. Scale bar: A, B: 5μm; A’-A’’’ and B’-B’’’:1 μm; H-K’: main image - 10μm, boxed inset - 1 μm.

Next, we examined IHC ribbon synapses at P12, the onset of hearing in mice (Mikaelian and Ruben, 1965). In P12 *Esrrg^+/+^* mice, abundant CtBP2-labelled ribbons were present in both mid-apical and mid-basal cochlear coils, accompanied by prominent clusters of GluA2-positive post-synaptic receptors (Fig. 3H’,J’). In contrast, although P12 *Esrrg*-cKO mice displayed clear CtBP2-positive ribbons, GluA2 signal intensity was markedly reduced (Fig. 3I’,K’). Strikingly, at 8 kHz, *Esrrg-*cKO mice exhibited a significant excess of CtBP2-positive ribbons compared with littermate controls, whereas a modest reduction was observed at 24 kHz (Fig. 3L). Consistent with adult analyses, P12 cKO mice also showed an increased number of orphan ribbons, particularly in the mid-apical coil (Fig.3N). Because CtBP2-positive ribbons were elevated at 8kHz at P12, we quantified colocalised synapses as a percentage of total CtBP2-positive ribbons per IHC (Fig. 3M). This analysis revealed that fewer paired synapses are already evident around the time of hearing onset (Fig. 3M).

### IHC synaptic vesicle fusion and basolateral membrane potassium currents are reduced in Esrrg-cKO mice

We then investigated whether the reduced ribbon synapses in IHCs of adult mice was associated with smaller synaptic exocytosis. Exocytosis in adult apical IHCs was estimated by measuring changes in cell membrane capacitance (*ΔC*_m_), following depolarising voltage steps that activate the Ca^2+^ current (*I_Ca_*). We found that the peak size of *I_Ca_* at -11 mV was not significantly different between the IHCs from adult *Esrrg-*cKO mice and *Esrrg^fl/fl^* controls (P = 0.0663, t-test, Fig. 4A-C). *ΔC*_m_ was investigated in response to a depolarising voltage step of -11 mV from 2 ms to 1.0 s in duration, which reveals the emptying of different populations of synaptic vesicle pools. When performing recordings at body temperature and using a physiological extracellular Ca^2+^ concentration (1.3 mM Ca^2+^), stimuli up to about 50 ms reveal the readily releasable pool (RRP), while longer steps cause the release of vesicles located further away from the Ca^2+^ channels (secondarily releasable pool: SRP) (Johnson et al., 2010, Johnson et al., 2017). We found the size of the isolated RRP (measured at 50 ms: Fig.4 D,F) and that of the total releasable vesicle pools (measured at 1 s: RRP + SRP: Fig. 4D,E) were both significantly reduced in *Esrrg-*cKO mice compared to *Esrrg^fl/fl^* controls (Fig. 4G,H). Collectively, these results support the reduced number of ribbons found in *Esrrg-*cKO mice.

**Figure 4.**
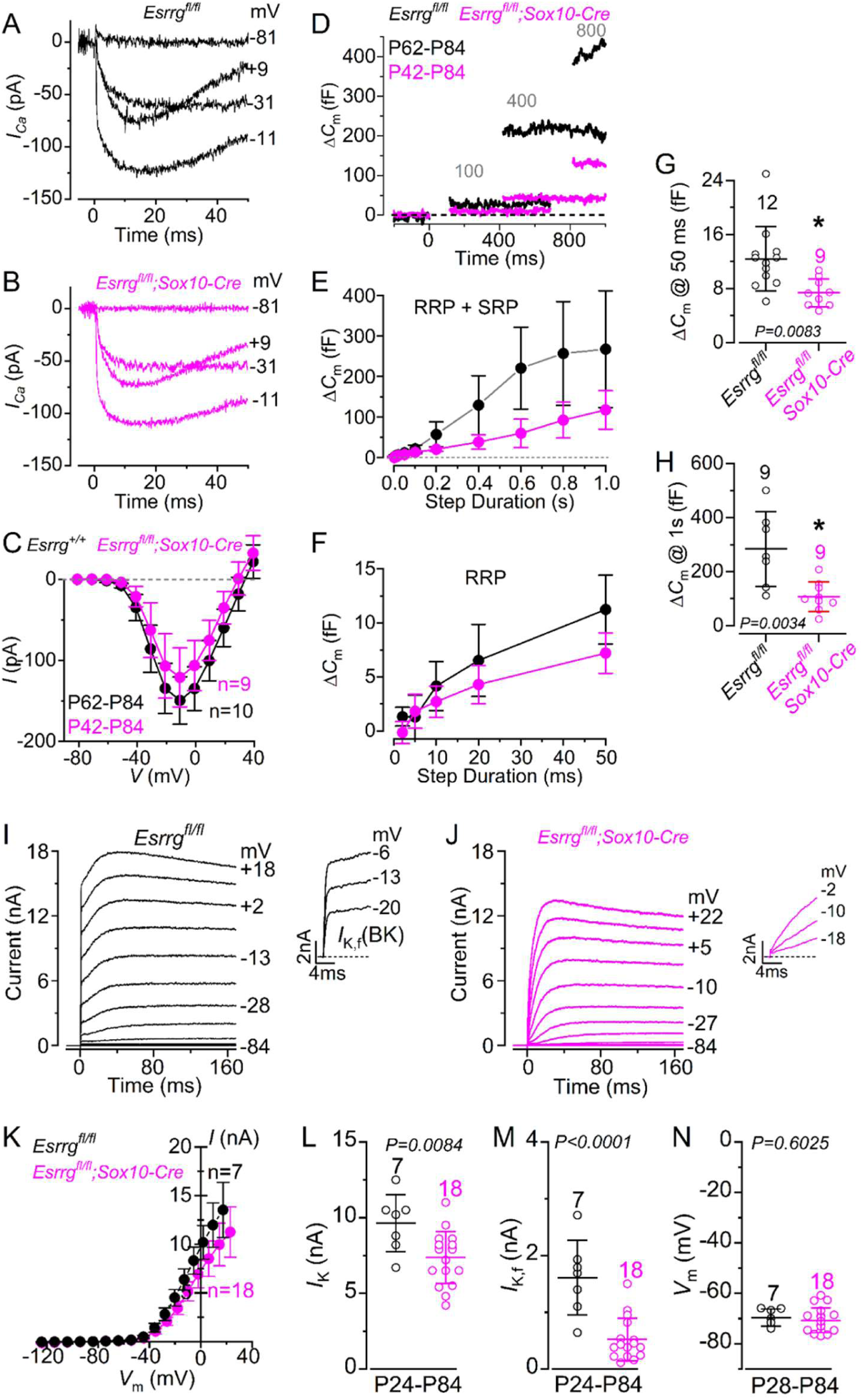
The biophysical properties of IHCs are impaired in *Esrrg*-cKO mice. **(A,B)** Example of calcium current (*I_Ca_*) recordings from IHCs of adult control **(A)** and *Esrrg^fl/fl^;Sox10-*Cre **(B)** mice obtained by applying depolarising voltage steps (10 mV increment) from -81 mV to more depolarized values. For clarity, only a few current recordings are shown. **(C)** Average peak Ca^2+^ current-voltage (*I_Ca_*-*V*_m_) curves obtained from IHCs of adult control (P62-P84) and *Esrrg^fl/fl^;Sox10-*Cre mice (P42-P84). Recordings were obtained as described in panels **A, B**. **(D)** Changes in cell membrane capacitance (*ΔC*_m_) obtained in response to depolarizations to -11 mV of varying duration (2 ms to 1.0 s; interstep interval was at least 11 s) from both genotypes. **(E)** Average *ΔC*_m_ recorded from IHCs of 9 controls (P62-P84) and 9 *Esrrg^fl/fl^;Sox10-*Cre mice (P42-P84) in response to voltage steps from 2 ms to 1.0 s (to -11 mV from the holding of -81 mV) showing the readily releasable pool (RRP: 2 ms to 50 ms) and the secondary releasable pool (SRP: 100 ms to 1.0 s). **(F)** Expanded view of the RRP (from 2 ms to 50 ms) from panel **E**. **(G,H)** Size of the total vesicle releasable pool measured in the IHCs at 50 ms (RRP, **G**) and at 1 s (RRP + SRP, **H**) from both genotypes. **(I, J)** Membrane K^+^ currents recorded from IHCs of control (**I**, P28, *Esrrg^fl/fl^*) and mutant (**J**, P24, *Esrrg^fl/fl^;Sox10-*Cre) mice. Currents were elicited by using depolarizing voltage steps, with a nominal increment of 10 mV, from a holding potential of −84 mV. Test potentials are shown next to some of the traces. The outward current is primarily carried by a delayed rectifier K^+^ current *I_K_* and a rapid activating Ca^2+^ activated K^+^ current *I_K,f_*, which is highlighted in isolation in the insets. Note that *I_K,f_* was largely reduced in the IHCs of *Esrrg^fl/fl^;Sox10-*Cre mice. **(K)** Steady-state current–voltage curves obtained from IHCs of *Esrrg^fl/fl^*(P24-P84) and *Esrrg^fl/fl^;Sox10-*Cre mice (P24-P84) mice. **(L,M)** Size of total steady-state outward K^+^ current **(L)** and the isolated *I_K,f_* **(M)** in the IHCs from both genotypes. Note that *I_K_* was measured at 0 mV, while the isolated *I_K,f_* was measured at 1 ms from the current onset and at -25 mV (Marcotti et al., 2003). **(N)** Resting membrane potential (*V_m_*) measured in IHCs from both genotypes. Data in **C, E, F-H, L-N** is plotted as mean values ± SD. **G,H,L-N** - open symbols: single cell values; the number of IHCs investigated is shown above the average data points from 7 controls and 9 *Esrrg^fl/fl^;Sox10-*Cre mice **(G,H)**; 4 controls and 8 *Esrrg^fl/fl^;Sox10-*Cre mice **(L-N)**.; statistical comparisons: unpaired t-test.

Considering the changes in synaptic activity, we investigated whether the basolateral membrane K^+^ currents in adult apical-coil IHCs were also disrupted in *Esrrg*-cKO mice. Membrane currents were elicited by applying depolarising voltage steps from the holding potential of –84 mV. Adult IHCs are known to express, in addition to the classical delayed rectifier K^+^ current (*I_k_*), a rapid Ca^2+^ activating K^+^ current, *I*_K,f_ (Kros et al., 1998, Marcotti et al., 2003, Marcotti et al., 2004). We found that both the total outward K^+^ current (*I*_K_) and the isolated *I*_K,f_ were significantly reduced in the IHCs of adult P24-P84 *Esrrg*-cKO mice compared to those recorded in cells from age-matched littermate controls (Fig. 4I-M). The strongly reduced or absence of ***I*_K,f_**, which is only expressed in adult IHCs, indicates that these cells were unable to fully mature, retaining a more pre-hearing phenotype. Despite the different amplitudes of K^+^ currents, the resting membrane potential measured in current clamp was comparable between the two genotypes (Fig. 4N). Collectively, these results indicate that *Esrrg* is necessary for IHCs to acquire their adult-like characteristics.

### *Esrrg* is required for normal cochlear wiring

Next, we investigated the auditory innervation using an antibody against neurofilament-heavy chain (NF-H) on wholemounts of the organ of Corti, since each type I SGN extends a single afferent fibre to an IHC ribbon (Meyer et al., 2009). To reveal myelinated fibres, we co-stained with CellMask™ orange (Wu et al., 2019) (Fig. 5A-H). In adult *Esrrg^+/+^* mice, the typical architecture of mature auditory innervation was observed with CellMask™ Orange staining the bulk of myelinated fibres as they traversed across the OSL, and NF-H immunolabelling revealing the short non-myelinated segments from the habenula perforata (HP) (Fig. 5A,C). We therefore presumed that most of these fibres were type I afferents, 90% of which are known to be myelinated (Nayagam et al., 2011). In contrast, in adult *Esrrg-*cKO mice, fibres labelled with NF-H rarely contained CellMask™ Orange along the full length of the neurite; in many fibres, the CellMask™ Orange signal stopped short of the HP, indicative of truncated myelination (Fig. 5B,D). Strikingly, in adult *Esrrg-*cKO mice, CellMask™ Orange staining revealed radial fibre bundles that were more sparsely distributed (Fig. 5D) and more closely resembled those characteristic of the immature cochlea (Katayama et al., 2009, Coate et al., 2012). Accordingly, the overall density of these fibres within the OSL was significantly reduced, particularly in the apex (Fig. 5I: CellMask™ Orange fluorescent intensity apex 5.60 ±0.94 x10^5^ versus 3.03 ±0.74 pixels x10^5^, *p*=0.0004 unpaired t-test; base 4.72 ±1.07 x10^5^ versus 3.26 ±0.35 pixels x10^5^, *p*=0.0112 unpaired t-test, in *Esrrg^+/+^*versus *Esrrg-*cKO mice, respectively). We then investigated whether the mutant phenotype originates early by examining the auditory innervation in the immature cochlea immediately after birth (Fig. 5F) and at P7, when many type I afferents are normally myelinated (Fig. 5H) (Long et al., 2018). Consistent with an early onset neuropathy, at P1, radial fibres were already grouped into more sparsely distributed bundles with large gaps evident between them (Fig. 5F). By P7, whereas CellMask™ Orange revealed fibres with abundant myelin ensheathment in *Esrrg*^+/+^ mice, truncated myelination was observed in *Esrrg*-cKO mice (Fig. 5G,H).

**Figure 5.**
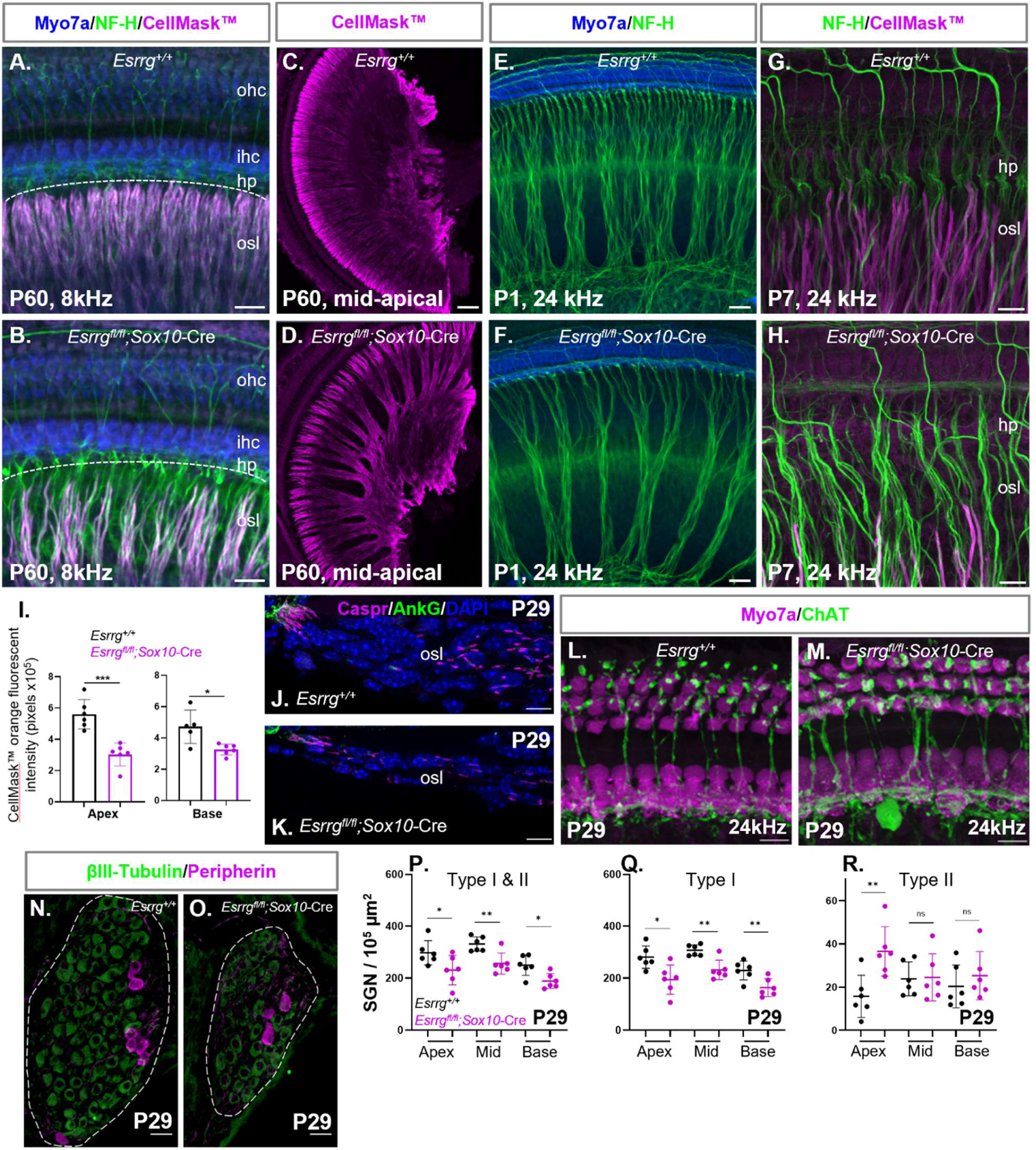
The auditory innervation is impaired in *Esrrg*-cKO mice from early development. Maximum intensity projections of confocal z-stacks of cochlear whole mount preparations from *Esrrg^+/+^* **(A,C,E** and **G)** and *Esrrg^fl/fl^;Sox10-*Cre **(B,D,F** and **H)** mice immunolabelled with antibodies to Myosin7a (Blue) and NF-H (Green), co-stained with the cell plasma membrane dye CellMask™ orange (Magenta) to reveal the myelinated fibres. *Esrrg^fl/fl^;Sox10-*Cre mice display defective myelination from early development **(H)** with reduced auditory fibres apparent at P1 **(F)**. **(I)** Abundance of myelinated fibres extracted from the fluorescent intensity of the CellMask™ orange signal at 8kHz and 24kHz from a 50µm by 59µm ROI in the OSL distal to the HP. ROI were acquired from 20x maximum intensity projections as shown in A, B. Data was acquired from 2-3 mice per genotype per sex. Maximum intensity projections of confocal z-stacks of apical coil cochlear cryosections from adult *Esrrg^+/+^* **(J)** and *Esrrg^fl/fl^;Sox10-*Cre **(K)** mice immunolabelled with antibodies to the heminodal proteins - Caspr (Magenta) and AnkG (Green). Maximum intensity projections of confocal z-stacks of cochlear whole mount preparations from adult *Esrrg^+/+^* **(L)** and *Esrrg^fl/fl^;Sox10-*Cre **(M)** mice immunolabelled with antibodies to the efferent innervation marker ChAT (green) and Myo7a (Magenta). Maximum intensity projections of confocal z-stacks of mid-coil cochlear cryosections from adult *Esrrg^+/+^* **(N)** and *Esrrg^fl/fl^;Sox10-*Cre **(O)** mice immunolabelled with antibodies to the pan-neuronal SGN marker βIII-Tubulin (Green) and the Type II marker Peripherin (Magenta). Note, Rosenthal’s canal (dashed lines) is smaller in *Esrrg^fl/fl^;Sox10-*Cre mice **(O)**. **(P-R)** SGN density at P29 in *Esrrg^fl/fl^;Sox10-*Cre mice compared to controls across basal, mid, and apical cochlear coils: **(P)** Type I & II, **(Q)** Type I and **(R)** Type II SGNs. Data was pooled from both male and female mice and acquired from 3 mice per genotype per sex. Data is plotted as mean values ± SD; *p<0.05; **p<0.01; ***p<0.001 unpaired t-test. Images represent data from 2-3 mice per genotype per sex. Scale bar: A,B: 20μm; C,D:100 μm; E,F: 20μm; G,H: 10μm; J,K: 10 μm; L,M: 10μm; N,O: 20 μm.

#### Disorganisation of first heminodes in *Esrrg-*cKO mice

The formation of first heminodes, the initial spike generators of type I SGNs, occurs in conjunction with axonal myelination during development (Kim and Rutherford, 2016, Smith et al., 2024). In rodents, spike generators form from a central position within the OSL around P5, then migrate towards the HP as they mature (Kim and Rutherford, 2016, Smith et al., 2024). Given the truncated myelination pattern in adult *Esrrg-*cKO mice, we next asked whether first heminodes were disrupted. Immunolabelling of cochlear cryosections with the paranodal junction protein Contactin-associated protein (Caspr) and the nodal protein Ankyrin-G (AnkG) (Smith et al., 2024) revealed the expected clustering of first heminodes in the HP region in adult *Esrrg^+/+^* mice (Fig.5J). In comparison, in *Esrrg-*cKO mice, first heminode density was greatly reduced, and Caspr-positive immunosignal was distributed irregularly throughout the OSL suggesting that in *Esrrg-*cKO mice, the IHC spike generator zone may fail to reach normal maturity (Fig.5K & Fig.S6).

#### Disorganised efferent innervation in Esrrg-cKO mice

In *Esrrg-*cKO mice, NF-H positive fibres traversing the tunnel of Corti appeared disorganised (Fig. 5B,F and H). Presumably, some of these are the unmyelinated Type II afferents. However, medial olivocochlear (MOC) efferent fibres that synapse on the OHCs also traverse the tunnel of Corti. Therefore, we immunolabelled adult organ of Corti wholemounts with an antibody against Choline acetyltransferase (ChAT) to examine the olivocochlear efferents (Sobkowicz and Emmerling, 1989) (Fig. 5L,M). This analysis revealed that MOC fibres traversing the tunnel of Corti and their OHC terminals, although present In *Esrrg-*cKO mice, appeared disorganised and sometimes thicker (Fig. 5M). Similarly, although the lateral olivocochlear (LOC) efferent fibres that extend along the inner spiral bundle were present, prominent large patches of immunofluorescence were often detected, suggestive of swollen LOC innervation (Fig. 5M). The gross morphology of the auditory innervation in *Esrrg-*cKO mice was then carefully examined in each sex, but no discernible differences were found within the limits of our analysis.

#### Loss of SGNs in *Esrrg-*cKO mice

Next, we investigated whether reduced numbers of SGNs could explain the sparsely populated auditory fibres in *Esrrg-*cKO mice. Immunolabelling of adult cochlear cryosections with the pan-neuronal marker βIII-Tubulin, revealed a reduced density of SGNs in *Esrrg-*cKO mice compared to control mice that reached significance in the apical (Fig. 5P; 297.55 SGN / 10^5^ µm^2^ ± 46.88 versus 230.62 SGN / 10^5^ µm^2^ ± 56.47, p=0.0495, t-test), mid (Fig. 5N-P; 331.22 SGN / 10^5^ µm^2^ ± 27.77 versus 256.30 SGN / 10^5^ µm^2^ ± 40.59, p=0.0039, t-test) and basal (Fig. 5P; 249.53 SGN / 10^5^ µm^2^ ± 38.98 versus 188.95 SGN / 10^5^ µm^2^ ± 28.52, p= 0.0118, t-test) cochlear coils.

We then used the Type II marker, Peripherin, to clarify if the reduced density of SGNs was due to a reduction in Type I or Type II neurons. This analysis revealed only the Type I neurons (βIII-Tubulin positive; Peripherin negative) were significantly reduced in each cochlear coil (Fig. 5N,O,Q; apical coil 280.63 SGN / 10^5^ µm^2^ ±43.11 versus 194.10 SGN / 10^5^ µm^2^ ±56.43, p=0.0137, t-test; mid coil 307.41 SGN / 10^5^ µm^2^ ±21.24 versus 231.77 SGN / 10^5^ µm^2^ ± 37.46, p=0.0016, t-test; and basal coil 229.19 SGN / 10^5^ µm^2^ ± 36.14 versus 163.66 SGN / 10^5^ µm^2^ ± 34.44, p=0.0092, t-test) in *Esrrg^+/+^*versus *Esrrg-*cKO mice respectively). The density of the Type II neurons (Peripherin positive) did not significantly differ except in the apical cochlear coil where they were significantly elevated. (Fig. 5R; 15.73 SGN / 10^5^ µm^2^ ±9.75 versus 36.52 SGN / 10^5^ µm^2^ ±11.41, p=0.0068, t-test, in *Esrrg^+/+^* versus *Esrrg-*cKO mice respectively). Strikingly, when examining the density of the SGNs, in addition to reduced numbers of Type I neurons, we found that the size of Rosenthal’s canal, which encases the SGNs, was significantly smaller in the mutant mice (Fig. S7). We then focused our investigations at P1 using the pan-neuronal marker, βIII-Tubulin, and found a similar trend, with significantly reduced numbers of SGNs encased in smaller canals in cKOs (Fig. S8B,C) suggesting that the reduction in SGN numbers in *Esrrg-*cKO mice occurs prior to birth. However, although the SGN density showed a trend towards a reduction in the cKO mice at P1, it did not reach significance (Fig. S8A), which we presume is partly due to the reduction in the size of Rosenthal’s canal masking the smaller number of neurons present. We then investigated the impact of sex on these findings, focusing on the density of Type I versus Type II SGNs in young adult *Esrrg*-cKO mice. We found that although each sex exhibited a trend toward reduced density of Type I SGNs, this only reached significance in females in the mid and basal cochlear coils, with female cKOs containing considerably fewer SGNs than male cKOs in the base. Similarly, the elevation in apical Type II SGN density only reached significance in females.

### Dysmyelination of auditory fibres in *Esrrg*-mutant mice

In mice, the ensheathment of auditory fibres with myelin lamellae commences early during postnatal development. By P12, around the time of hearing onset, the myelination of the majority of the type I afferents has successfully extended to the HP where it terminates at the spike generator zone (Long et al., 2018). Consequently, we reasoned that in adult *Esrrg-*cKO mice, where the auditory myelination is truncated, the myelin ultrastructure may exhibit features more typical of an immature phenotype. To better understand the myelin defect, we used transmission electron microscopy (TEM) on apical thin sections of the organ of Corti (Fig. 6). Consistent with our CellMask™ Orange staining (Fig. 5B,D,H,I), TEM thin sections showed that the density of the myelinated auditory fibres as they traversed the OSL was reduced in *Esrrg-*cKO mice compared to *Esrrg^+/+^*controls (Fig. 6A,B). Further, the thickness of the myelinated fibres in *Esrrg-*cKO mice often appeared thinner (Fig. 6D magenta arrows) with fewer mitochondria present (Fig. 6D magenta asterisks). To examine the thickness of the myelin sheath in more detail we used TEM to measure the width of the myelin sheath (Fig. 6E). This analysis revealed a significant reduction in myelin sheath thickness in *Esrrg-*cKO mice compared to control mice (Fig. 6F: 254.80nm ±73.02 versus 307.90nm ±65.86, respectively *p*<0.0001, t-test) concomitant with a significant reduction in the number of individual myelin lamellae layers (Fig. 6G: n= 20 ±5 versus 23 ±4 lamella, respectively *p*=0.0188, Mann-Whitney U test). These findings, together with our immunolabelling investigations (Fig. 5B,D,H), suggest that *Esrrg* is required for the myelin sheath of peripheral auditory fibres to reach full maturity. Stratifying these results by sex revealed similar results in both males and females with mutant mice displaying thinner myelin sheaths with fewer lamellae; however, these differences only reached significance in females (Fig.S10 A,B).

**Figure 6:**
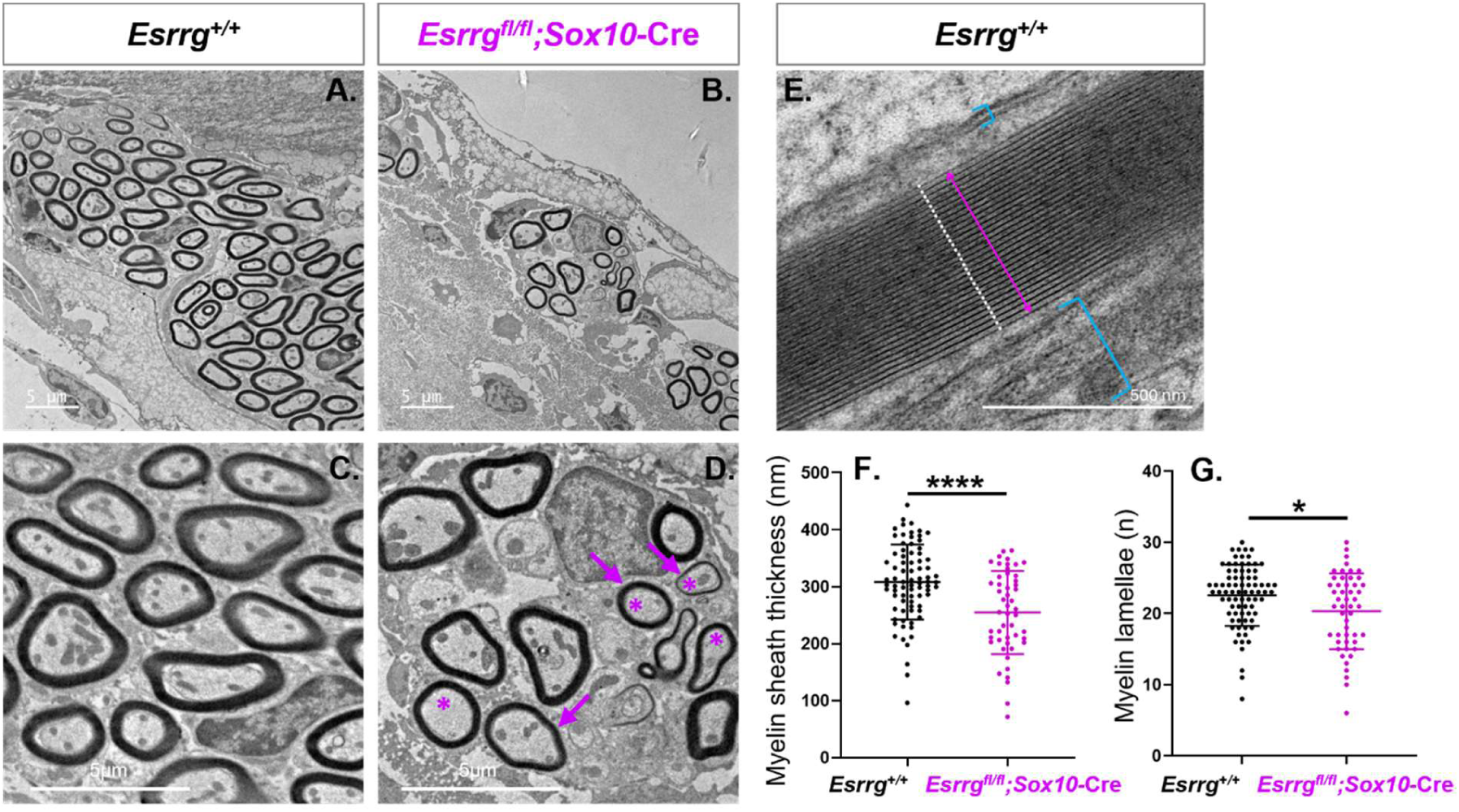
Aberrant myelin ultrastructure in *Esrrg*-cKO mice. TEM thin sections of apical auditory nerve fibres in the OSL from adult (P57-P65) *Esrrg^+/+^* **(A,C,)** and *Esrrg^fl/fl^;Sox10-*Cre **(B,D)** mice show reduced myelinated fibres in cKOs. **(D)** Note the thinner fibres in *Esrrg^fl/fl^;Sox10-*Cre mice mice (magenta arrows) with fewer mitochondria (magenta asterisks). **(E)** Representative TEM thin section from an *Esrrg^+/+^* mouse to show how myelin sheath thickness and individual lamellae were quantified. Myelin sheath lamellae were easily distinguished by their uniformity and density (E: white dots) and were manually counted using Fiji software, V2.15.0. The thickness of the myelin sheath was measured by measuring the width of the myelin perpendicular to the first and last lamella (E: magenta arrow); the axolemma and neurilemma were excluded (E: small and large aqua brackets, respectively). Myelin sheath thickness **(F)** and number of myelin lamellae **(G)** in *Esrrg^+/+^*(n=9; 4 females, 5 males) versus *Esrrg^fl/fl^;Sox10-*Cre mice (n=7; 4 females, 3 males) mice. Metrics were obtained from at least 8 apical auditory nerve fibres per sample. Statistical comparisons: unpaired t-test (\*\*\*\**p*<0.0001) or Mann–Whitney U test (\**p*<0.05) depending on the normality of the data.

### Gene expression profiling in *Esrrg*-cKO mice

To identify the differentially expressed genes (DEGs) in *Esrrg*-cKO mice that could explain the early-onset auditory defect, we performed gene expression profiling by RNA-Seq analysis on whole cochlear tissue from female *Esrrg^fl/fl^;Sox10-Cre* mice and *Esrrg^fl/fl^* littermate controls at P1 (Fig.7). A total of 179 DEG met our inclusion criteria and were significantly dysregulated (Fig.7A,B; Table S3). Consistent with the role of *Esrrg* as a consecutively active transcription factor (Huss et al., 2015), the majority of the DEGs (157) were significantly downregulated in *Esrrg*-cKO mice, while only 22 exhibited significant upregulation (Fig.7A,B). Since the density of the SGNs was reduced in *Esrrg*-cKO mice and our data suggested this occurs prior to birth (Fig. 5P-R; S8), changes in gene expression such as these could reflect indirect changes on the cellular level in addition to specific changes within existing cells. Therefore, to determine which DEGs might be direct targets of *Esrrg* we performed *in-silico* analysis with iRegulon predictive software using a motif for the estrogen-related receptor response element, ERRE (Dufour et al., 2007, Sadasivam et al., 2024). This analysis revealed 65 DEGs (36%) contained putative binding sites for *Esrrg* suggesting they could be direct targets (Fig.7C). We also compared the expression of 33 common housekeeping genes between genotypes, including the pan-neuronal marker βIII-Tubulin, but, apart from with housekeeping gene *Tfrc,* we found no significant differences in expression (Table: S4). Interestingly, *Tfrc,* a mitochondrial regulator (Senyilmaz et al., 2015), was also a hit for one of our direct putative targets (Fig. 7C). Subsequent qPCR analysis of a subset of putative target genes confirmed results consistent with our bulk RNA-Seq analysis (Fig. S11).

**Figure 7.**
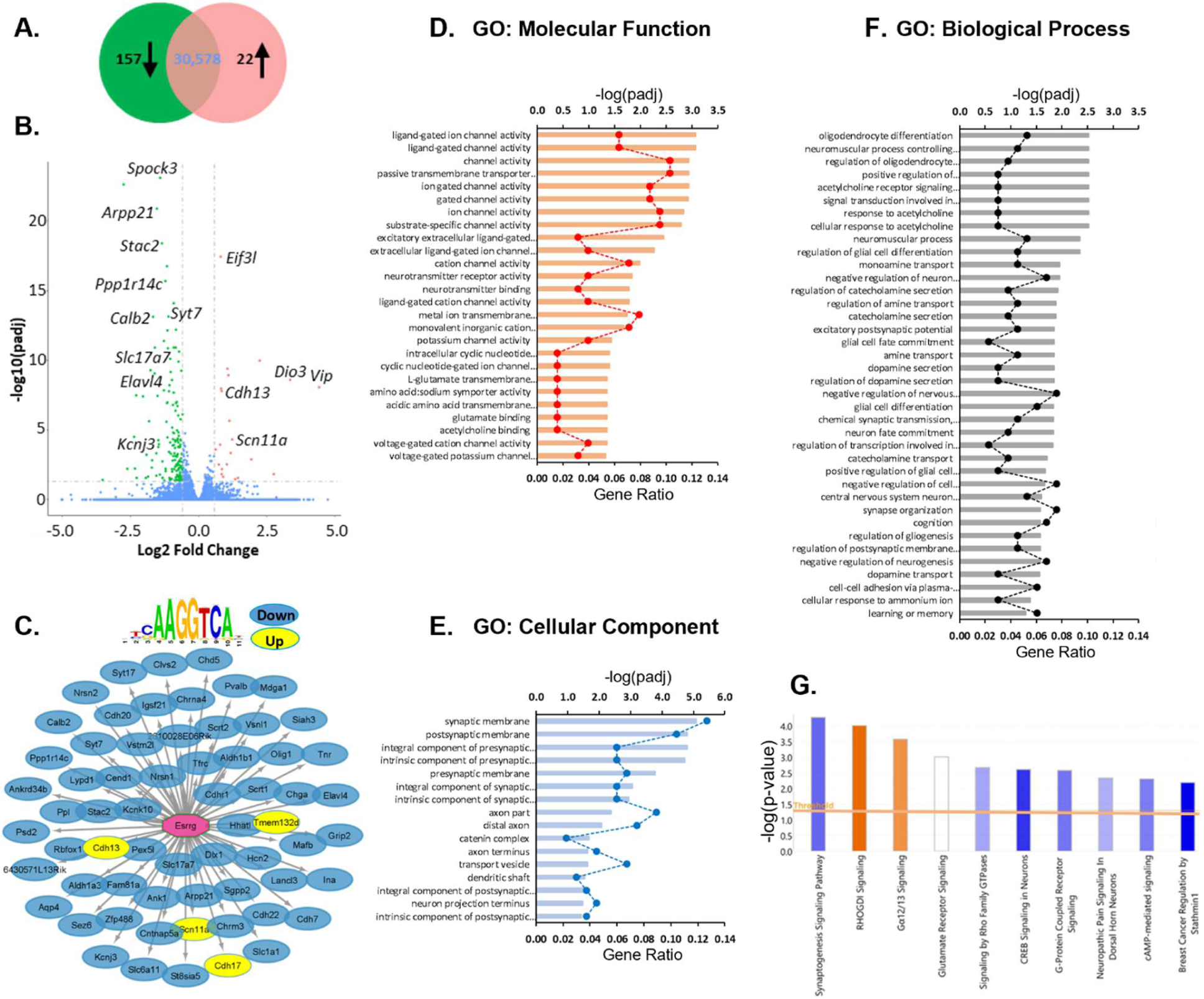
RNA-Seq and pathway analysis in *Esrrg*-cKO mice. Results of cochlear gene expression profiling from female *Esrrg^fl/fl^* (n=5) and *Esrrg^fl/fl^;Sox10-*Cre (n=5) mice at P1. Only DEGs that met our inclusion criteria (log2FoldChange ± ≥ 0.585; padj < 0.05) were included in subsequent analysis. **(A)** Venn diagram and **(B)** Volcano plot to illustrate the distribution of the DEGs in *Esrrg^fl/fl^;Sox10-*Cre mice. **(B)** Each point represents a single DEG; down-regulated (Green), up-regulated (Orange); DEGs that did not meet our inclusion criteria are shown in Blue. **(C)** DEGs with a putative binding motif for the estrogen-related receptor response element (ERRE). Data was generated using iRegulon predictive software. DEGs that were down-regulated in *Esrrg^fl/fl^;Sox10-*Cre mice are shown in blue, up-regulated DEGs are shown in yellow. **(D-F)** GO enrichment analysis of DEGs ranked by *padj* <0.05; **(D)** GO: Molecular Function, **(E)** GO: Cellular Component, **(F)** GO: Biological Process. Dashed line: Gene ratio for each GO category (number of DEGs annotated per GO Term / Total number of DEGs). **(G)** Ingenuity pathway analysis to show the top 10 *Canonical Signaling Pathways* associated with the DEGs in *Esrrg^fl/fl^;Sox10-*Cre mice. Pathways are ranked according to p-value (*P* <0.05 (-log *p*=1.3), Fisher’s Exact Test, right-tailed). Significance (threshold) is denoted by the orange line. Z-scores were used to denote whether a pathway is predicted to be inhibited or activated in the DEGs. Blue: negative z-score (inhibitory); orange: positive z-score (activation) – the stronger the shading, the higher the value of the z-score.

The most significant DEG on our down list was *Spock3* encoding osteonectin, a calcium-binding glycoprotein that is strongly induced in cochlear supporting cells prior to hearing onset (Mothe and Brown, 2001). Notably, *Gria2,* which encodes the GluA2 protein and was significantly reduced in our immunolabelling studies at the IHC ribbon synapse (Fig.3), was also downregulated at the transcript level. Crucially, some of our putative downstream targets of *Esrrg* in the cochlea corroborated findings from previous studies in neuronal tissue (Pei et al., 2015, Fox et al., 2025), including *Stac2*, the 4^th^ most significant downregulated DEG, which encodes an adaptor protein that suppresses Ca^2+^-dependent inactivation of neuronal L-type Ca^2+^ channels in the brain (Polster et al., 2018), and *Slc6a11*, a sodium-dependent transporter that imports the inhibitory neurotransmitter GABA (Scimemi, 2014). We also found that two molecular markers of the Type I SGN subtypes, *Calb2,* which is highly expressed in subtype 1a neurons, and *Lypd1,* which is highly expressed in subtype 1c neurons (Shrestha et al., 2018), were putative targets of *Esrrg* on our down list (Fig.7C). Additional putative targets that were downregulated included several RNA-binding proteins - *Arpp21,* involved in dendritic branching in cortical neurons (Rehfeld et al., 2018); *Rbfox1,* associated with the maturation of neuronal circuits in the brain (Prashad and Gopal, 2021); and *Elavl4,* widely linked to neurogenesis and neuronal maturation (Mulligan and Bicknell, 2023, Wutikeli et al., 2025). *Mdga1*, involved in synaptic structural integrity (Lee et al., 2023) and *Psd2* implicated in axon growth and synaptogenesis (Wu et al., 2020) were also listed. Also, of interest, was *Mafb,* involved in the refinement of the Type I SGN subtypes and their associated synaptic connections (Bastille et al., 2025), and *Hcn2*, a hyperpolarisation-activated cation channel that is broadly expressed in the cochlear sensory and neuronal tissue (Luque et al., 2021) and a key modulator of myelin sheath length in the central nervous system (Swire et al., 2021).

The most significant upregulated DEG was *Eif3l*, encoding eukaryotic translation initiation factor 3 subunit L involved in the initiation of protein synthesis, particularly in neurons (Fig.7B) (Gomes-Duarte et al., 2018). Cadherin genes, *Cdh13* and *Cdh17,* and the voltage-gated sodium channel subunit *Scn11a* (*Nav1.9*), implicated in IHC synaptic function (Zu et al., 2021) were also upregulated and hits for direct putative targets (Fig.7C). Unexpectedly, also on our Up list was *Esrrg* (log2 fold-change: 0.623632; padj: 0.00026); albeit close to the cut-off for our inclusion criteria. Since this knockout strategy selectively disrupts the critical exon, presumably this is a consequence of the upregulation of the 3’ flanking exons in response to the lack of the critical exon (Davuluri et al., 2008) (Fig.1C).

To probe our DEGs in more detail we used Gene Ontology (GO) enrichment analysis (Fig.7D-F). GO terms associated with *Molecular Function* that were significantly enriched amongst the downregulated genes included *ligand-gated ion channel activity* (eg: *Slc17a7*, *Chrna4, Kcnj3, Glra2*, *Gria2*, *Hcn2)*, *neurotransmitter receptor activity* (eg: *Htr7, Chrna4, Glra2, Gria2,* Chrm3), potassium *channel activity* (eg: *Kcnj3, Kcnk10, Kcnt1), L-glutamate transmembrane transporter activity* (eg: *Slc17a7, Slc1a1), amino acid sodium symporter activity (*eg*: Slc6a11, Slc1a1)* and *acetylcholine binding* (eg: *Chrna4, Chrm3*) (Fig.7D). Significant GO terms associated with *Cellular Components* were mainly involved with neurotransmission including *synaptic membrane* (eg: *Syt7, Calb2, Lrrtm3, Htr7, Grm7, Chrna4, Ank1, Slc6a11 Glra2, Gria2, Grip2, Igsf21), axon part* (eg: Syt7, *Elavl4, Ank1)* and *transport vesicle (eg: Syt7, Slc17a7, Syt17, Nrsn2, Nrsn1*) (Fig.7E). Similar GO terms associated with *Biological Process* included *excitatory postsynaptic potential* (eg: *Sez6, Slc17a7, Chrna4, Cbln1, Glra2, P2rx3)* and *synapse organization* (eg: *Sez6, Mdga1, Lrrtm3, Sema3e, Tnr, Igsf21, Slit1)* (Fig.7F). Additionally, we also found GO terms linked with *oligodendrocyte differentiation* (eg: *Olig1, Olig2, Dlx1, Aspa, Pax6, Zfp488, Nkx2-*2,) were enriched. This led us to examine the expression of common glia markers within our DEGs. However, transcripts for the neural crest transcription factor, *Sox10*, a regulator of peripheral glial development that is also expressed in oligodendrocyte precursor cells (Kuhlbrodt et al., 1998, Watanabe et al., 2000, Britsch et al., 2001) and *Sox2*, an early marker of cochlear satellite and Schwann cell glia (Smith et al., 2021) did not significantly differ between genotypes (S5: Table). No GO terms in any subcategory were significantly enriched amongst the upregulated DEGs in *Esrrg*-cKO mice apart from GO: *cellular component;* the *catenin complex* was significantly associated with two cadherin genes, *Cdh13 and Cdh17*.

Next, we used Ingenuity Pathway Analysis to identify the canonical signaling pathways associated with the DEGs (Fig.7G). The most significant pathway was the *Synaptogenesis Signaling Pathway* (involving *Gria2*, *Grm7*, *SYT17*, *SYT7* and members of the Cadherin gene family) which was predicted to be inhibited in *Esrrg-*mutant mice (-log p-value = 4.27; z-score = - 1.897); hence, suggesting that *Esrrg* normally plays a positive role in synaptogenesis. We also compared our DEGs to a manually curated list of 904 genes implicated in hearing from human and/or mouse studies (Lewis and Steel, 2025) and found that 7% of all DEGs were known deafness genes; of these, 42% overlapped with our iRegulon analysis and contained a putative *Esrrg-*binding site (Table: S6).

## Discussion

### Hearing loss in *Esrrg-*cKO mice is characteristic of auditory neuropathy

Our interest in *ESRRG* stemmed from our previous discovery linking this gene to susceptibility to ARHL in women (Nolan et al., 2013). Here, by generating a new *Esrrg* cKO mouse, we provide the first mechanistic insight into the role of *Esrrg* in the cochlea. We show that hearing loss in *Esrrg*-cKO mice is consistent with auditory neuropathy and provide evidence that *Esrrg* is a crucial molecular driver of auditory innervation and the functional maturation of the cochlea. These findings add to the growing evidence that the ERR family is essential for hearing (Chen and Nathans, 2007, Collin et al., 2008, Ben Said et al., 2011, Lee et al., 2011, Safka Brozkova et al., 2012, Nolan et al., 2013, Bhatt et al., 2016, Schilit et al., 2016, Khan et al., 2019).

Mice are not born hearing. Instead, the early postnatal murine cochlea must undergo a tightly coordinated series of developmental processes that culminate in hearing onset around P12 (Mikaelian and Ruben, 1965, Coate et al., 2019). Compared with *Esrrg^+/+^* littermates*, Esrrg*-cKO mice exhibit significant hearing loss by 2 weeks of age indicating that ESRRG is required for normal hearing development. At this stage, *Esrrg*-cKO mice also display abnormal tone-evoked ABR waveform morphology with reduced wave 1 amplitudes. We attribute this, in part, to early-onset myelination and synaptic defects, as afferent fibre myelination lagged markedly behind that of littermate controls at P7, and by P12, IHC ribbon synapses were reduced. These features closely resemble those reported in *Bace1^-/-^* mice, which develop auditory impairment due to delayed peripheral myelination combined with IHC ribbon synaptic defects (Dierich et al., 2019). Further, at P12, the GluA2 receptors detected at the post-synaptic density were poorly defined in *Esrrg-*cKO mice but not in littermate controls. Instead, they were more characteristic of the diffuse GluA2 morphology typically observed much earlier in auditory development, during the first postnatal week (Liberman and Liberman, 2016). Consequently, we speculate that although some GluA2 receptors form in *Esrrg-*cKO mice, they fail to progress in a timely manner along their normal developmental trajectory.

In mice, it is well documented that auditory innervation and IHC ribbon synapses continue to mature in the weeks following hearing onset, along with the molecular identities of the SGN Type I subtypes (Ia, Ib and Ic—widely thought to correspond to physiological subtypes), reaching their mature form around one month (Bulankina and Moser, 2012, Liberman and Liberman, 2016, Petitpre et al., 2018, Shrestha et al., 2018, Sun et al., 2018). In *Esrrg*-cKO mice, although ABR thresholds showed some improvement by young adulthood, they remained significantly elevated, and the reduction in wave 1 amplitude observed at 2 weeks did not recover, which we reasoned is partly explained by the persistent synaptic and myelination defects.

Strikingly, Rosenthal’s canal was smaller and surrounded fewer SGNs in *Esrrg-*cKO mice at P1. This anomaly continued into young adulthood, with the density of Type I SGNs significantly reduced in *Esrrg-*cKO mice across each cochlear coil. These results agree with a previous study in striatal spiny projection neurons showing that developmental deletion of *Esrrg* leads to reduced neuronal density (Fox et al., 2025). This reduction in SGN number also explains why the radial auditory innervation at P1 and in young adulthood appears sparse, as fewer Type I neurons are available to extend neurites. It further helps account for the reduced number of paired ribbon synapses detected at P12 and in adulthood, as well as the decreased first heminode density in the spike generator zone. In contrast to the reduction in Type I SGNs, Type II SGN density appeared to increase in the apical cochlear coil of *Esrrg-*cKO mice, but not in the mid or basal coils. Presumably, this increased density of apical Type II SGNs, at the expense of fewer Type I SGNs innervating apical IHCs, indirectly contributes to the elevated ABR thresholds at low frequencies.

There was no obvious loss or ultrastructural abnormalities of the sensory hair cells in *Esrrg*-cKO mice, and DPOAE recordings were normal. These findings suggests that ESRRG is not required for sensory hair cell survival, at least through young adulthood, and that defective outer hair cell function does not account for the auditory deficit. The auditory ossicles also appeared normal indicating that abnormal middle ear function was unlikely to explain the raised ABR thresholds, particularly in the lower frequencies. However, we cannot exclude a subtle role for *Esrrg* in OHC function and cochlear amplifier activity, as the ChAT-labelled MOC efferent fibres that synapse on the OHCs often appeared thicker and disorganised. Similarly, the LOC efferent fibres, which synapse with the Type I afferents, often appeared swollen.

Despite the role of ESRRG in energy metabolism and in regulating potassium ion homeostasis in organs that, like the cochlea, have high metabolic demand (Alaynick et al., 2007, Dufour et al., 2007, Alaynick et al., 2010), the EP was normal in young adult *Esrrg-*cKO mice. This suggests that reduced activity of the stria vascularis does not contribute to the auditory phenotype. The closely related nuclear receptor ESRRB is crucial for endolymph production by the stria vascularis through regulation of multiple ion channel and transporter genes (Chen and Nathans, 2007). Therefore, it is conceivable that either ESRRG does not play a major role in EP generation, or ESRRB compensates for the loss of ESRRG.

Collectively, this body of evidence, particularly, the elevated ABR thresholds in the presence of normal DPOAE and EP recordings, together with synaptic and neuronal defects in the absence of sensory hair cell loss, strongly indicates that the prime origin of the hearing loss in *Esrrg*-cKO mice is auditory neuropathy (Shearer and Hansen, 2019).

Notably, given our previous study linking *ESRRG* with ARHL susceptibility in women, we included sex as a biological variable into our investigations. However, we did not identify a clear mechanism by which *ESRRG* mediates sex-specific effects on hearing. *Esrrg*-cKO mice of both sexes showed comparable elevations in ABR thresholds and similar reductions in wave 1 amplitude with prolonged latency.

### Functional maturation of key components of the auditory machinery is delayed in *Esrrg-*cKO mice

The auditory neuropathy in *Esrrg-*cKO mice affects multiple aspects of auditory innervation, with key components of the auditory machinery failing to reach functional maturity. Young adult *Esrrg*-cKO mice exhibit spike generator zones that are sparsely populated with first heminodes. They also show aberrant spacing between the first heminode and the first node of Ranvier and display ectopically positioned heminodes (Caspr-positive) within the intervening region of the OSL (Fig. 5K; S6). We propose that these findings indicate a failure of first heminodes to migrate along the peripheral afferents to their mature position at the spike generator zone, because this process is normally largely complete by the end of the second postnatal week (Kim and Rutherford, 2016, Smith et al., 2024). In addition, myelination of the peripheral auditory fibres, which normally reach the HP around hearing onset at P12 (Long et al., 2018), remained delayed in adult *Esrrg*-cKO mice at 2 months, and the myelin sheaths were thinner—features consistent with delayed maturation (Dierich et al., 2019). During auditory development, heminodes assemble along the advancing edge of myelin (Romand and Romand, 1985, Kim and Rutherford, 2016). Thus, the truncated myelination in the OSL parallels the dispersed localisation of first heminodes in this region and their reduced clustering at the spike generator zone. Cochlear nerve myelination and first heminode alignment are essential for propagating the IHC post-synaptic response by encoding sound-driven spikes (Long et al., 2018), and their disruption is linked to reduced wave 1 amplitude with prolonged latency (Wan and Corfas, 2017, Cassinotti et al., 2024). We therefore propose that myelin and heminode defects are largely responsible for the reduced wave 1 amplitude with prolonged latency in adult *Esrrg*-cKO mice.

Although we did not find a functional impact on the OHCs, we found that the biophysical properties of the IHCs were disrupted. In addition to the uncoupling at the IHC ribbon synapse, synaptic vesicle fusion (exocytosis) was significantly reduced and both the IHC *I_k_* and *I_k,f_* outward potassium currents in our adult Esrrg-cKO mice were markedly decreased appearing more comparable to those observed in postnatal mice prior to hearing onset (Marcotti et al., 2003). This suggests that *Esrrg* is required for IHCs to acquire their mature characteristics essential for normal pre-synaptic function. Our bulk RNA-seq and in-silico analyses suggest that this may be a direct effect, given the high proportion of DEGs containing an ERRE. However, we cannot exclude an indirect mechanism, since the anomalies observed in efferent innervation may have contributed to the IHC synaptic defect and, consequently, the lack of IHC functional maturation (Johnson et al., 2013, Yin et al., 2014).

### *Esrrg* is a molecular driver of auditory innervation maturation

As a transcription factor, *Esrrg* is ideally positioned to orchestrate gene networks, ensuring that the normal trajectory of auditory innervation development and maturation proceeds in a timely and coordinated manner. In the heart, *Esrrg* regulates genes essential for nearly all aspects of cardiac maturation; without ESRRG, cardiomyocytes remain functionally immature (Alaynick et al., 2007, Dufour et al., 2007, Alaynick et al., 2010, Sakamoto et al., 2020). Similarly, in the absence of ESRRG, pancreatic β-cells fail to metabolise glucose and mature into insulin-secreting cells (Yoshihara et al., 2016), while stomach parietal cells fail to mature into functional gastric acid-secreting cells (Alaynick et al., 2010, Adkins-Threats et al., 2024). Together with similar findings in kidney (Berry et al., 2011) and muscle (Murray et al., 2013), these studies support a model in which *Esrrg* functions as a master regulator of functional maturation across diverse, highly metabolic tissues to drive normal physiological function. We argue that the data described herein, including our bulk RNA-Seq and subsequent *in-silico* analysis suggests that *Esrrg* plays a similar role in the cochlea.

A broad range of putative target genes of *Esrrg* were identified, some of which are essential for normal cochlear innervation, whereas others have been linked to neuronal function but not previously implicated in the cochlea. Interestingly, in agreement with the growing role of RNA-binding proteins in neurodevelopment (Parra and Johnston, 2022), several genes encoding RNA-binding proteins (*Arpp21, Rbfox1, Elavl4*) (Rehfeld et al., 2018, Prashad and Gopal, 2021, Wutikeli et al., 2025) were among our putative *Esrrg* targets suggesting that regulation of RNA-binding proteins may be one mechanism through which *Esrrg* functions as a molecular driver in the cochlea.

Although the EP was normal in *Esrrg*-cKO mice, a subset of *Esrrg* targets included genes encoding potassium channels, as well as genes involved in calcium ion homeostasis and transporter activity, consistent with previous studies (Dufour et al., 2007, Alaynick et al., 2010). Notably, some downregulated putative *Esrrg* targets are involved in glutamate transmembrane transport, raising the notion that defective glutamate signalling contributes to the auditory deficit. The reduced number of IHC ribbon synapses, together with the IHC exocytosis defect supports this concept, since less glutamate is likely to be available for neurotransmission.

Further, consistent with previous reports linking *Esrrg* to the regulation of synaptic genes (Fox et al., 2022), several of our putative *Esrrg* targets are important for synapse formation and structural integrity including *Psd2,* and *Elavl4.* In line with this, the top canonical signalling pathway associated with our DEGs was the *Synaptogenesis Signalling Pathway.* Interestingly, both *Psd2,* and *Elavl4* have been reported to be upregulated during synaptogenesis and the regeneration of afferent neurites to hair cells in *ex-vivo* neonatal cochlear explants (Wu et al., 2020). Although some of the disruption at the IHC ribbon synapse can be attributed to a direct consequence of fewer SGNs and hence, fewer peripheral neurites, the above findings indicate that *Esrrg* has a direct role in the regulation of synaptogenesis in the cochlea. Consequently, it is plausible that disruption at the IHC ribbon synapse is one mechanism whereby genetic variation in *ESRRG* leads to ARHL in humans, particularly in women. This raises the possibility that targeted therapies focused on maintaining ESRRG may also preserve hearing.

## Materials and Methods

### Ethics statement

Mouse studies were conducted in accordance with the U.K. Animals (Scientific Procedures) Act of 1986 (ASAP) under U.K. Home Office licenses. All studies were approved by King’s College London and University of Sheffield Animal Welfare and Ethical Review Bodies. Mice had free access to food and water and a 12-hour light/dark cycle. Adult mice were killed by cervical dislocation (primary) using secondary methods as appropriate under these licenses. Postnatal mice less than P9 were culled by decapitation.

### Generation of Esrrg-*cKO* mice

Embryos from mice carrying a floxed allele for *Esrrg*, B6J.129S2-*Esrrg*^tm1.1Ics^/Ics; MGI ref: 6476827, (hereafter referred to as *Esrrg^fl^)* were sourced from the Institut Clinique de la Souris (Strasbourg, France). *Esrrg^fl/fl^* mice contain LoxP sites flanking critical exon 2, which encodes a transcriptional activation domain and part of the DNA binding domain (Misawa and Inoue, 2015) (Fig.1A). To disrupt *Esrrg* in the inner ear, *Esrrg^fl/fl^* mice were crossed with mice carrying the *Sox10-Cre* transgene (B6;CBA-*Tg(Sox10-Cre)1Wdr*/J; MGI ref: 3586900; provided by Prof. William Richardson, University College London) to generate *Esrrg-*cKO mice (*Esrrg^fl/fl^;Sox10-Cre* mice) on a mixed genetic background (Fig.1A). Sox10 is expressed around embryonic day 9.5 in the developing otic vesicle and in craniofacial neural crest-derived cells (Southard-Smith et al., 1998, Watanabe et al., 2000); consequently, *Sox10-Cre* selectively targets *Esrrg* throughout the inner ear. To avoid aberrant homologous recombination, the *Sox10-Cre* transgene was transmitted via the maternal germline as previously described (Crispino et al., 2011). Mice were born at standard Mendelian ratios, and littermates were used as controls in all experiments: either homozygous wildtype at the *Esrrg^fl^* locus and positive for *Sox10-Cre* (*Esrrg^+/+^*) or homozygous for the *Esrrg^fl^* allele and negative for *Sox10-Cre* (*Esrrg^fl/fl^*). In some experiments, *Esrrg^+/+^* and *Esrrg^fl/fl^* mice were combined as controls and, for simplicity, are hereafter referred to collectively as *Esrrg*^+/+^. Both male and female mice were used; sex breakdown and sample sizes are provided in the figure legends.

### Genotyping

DNA was extracted from pinna tissue, and genotyping was performed using the following primers as previously described (Murray et al., 2013): Lf87 (5’-CCCTTATGCTGATTACCTTCTTGTA-3’), Lr88 (5’-CAACAATGTAGACACAAAGACATGG-3’), Ef610 (5’-GTTTTAAAGGCCCTTGGTGATCTCGC-3’) and Er612 (5’-CTGCAACCCTTGGACTGCCAGAAC-3’) (Fig.1A). PCRs were run with Lf87 and Lr88, which span the 3’ LoxP site and generate 161bp or 208bp bands, and with Ef610 and Er612, which span the 5’ LoxP site and generate 149bp or 288bp bands, to distinguish between the native and floxed loci, respectively. An additional PCR was run with Lf87 and Er612, which generates a 232bp band in the presence of the recombined allele only. Confirmation of the *Sox10-Cre* transgene was detected using *Sox10-Cre-*specific primers, Sox10Cre_F_23316 (5’-CACCTAGGGTCTGGCATGTG-3’) and Sox10Cre-R_oIMR9074 (5’-AGGCAAATTTTGGTGTACGG-3’), which generate a 300bp band.

### Anaesthesia for in vivo physiology

For ABR recordings mice were anaesthetised with ketamine (100 mg/kg, i.p., Ketaset, Fort Dodge Animal Health) and xylazine (10 mg/kg, i.p., Rompun, Bayer Animal Health); recovery, where applicable, was promoted using atipamezole (1 mg/kg, i.p., Antisedan, Pfizer). For DPOAE recordings and EP measurements, mice were anaesthetised with intra-peritoneal 0.1 ml / 10 g urethane (20% w/v solution of urethane in water).

#### ABR Recordings

Brainstem auditory evoked potentials were measured as previously described (Ingham, 2019). Anaesthetised mice were placed in a sound-attenuating chamber and subcutaneous needle electrodes were inserted on the vertex (active) and overlying the left (reference) and right (ground) bullae. Responses were recorded to free-field calibrated broadband click stimuli (10 µs duration) and tone pips (5 ms duration, 1 ms onset/offset ramp) at 3, 6, 8, 12, 18, 24 and 30 kHz over 0-95 dB SPL (in 5 dB steps). Stimuli were generated via custom software on Tucker Davis Technologies (TDT) System 3 hardware (RP2.1 processors) and presented via a CTS Type 341 sound transducer. Evoked responses were amplified, digitised, and bandpass filtered between 300-3000 Hz, using custom software and TDT hardware (RA16 & RP2.1 processors, RA4LI low impedance headstage, RA4PA preamplifier). Thresholds of ABRs were defined as the lowest stimulus level to evoke a visually identifiable waveform. Evoked ABR waveforms were analysed offline and the latency and amplitude of ABR wave 1 plotted as a function of sound pressure level.

#### DPOAE recordings

Measurements of the 2f1-f2 DPOAE component were made as previously described (Buniello et al., 2016, Ingham et al., 2021) from mice positioned on a heating blanket inside a sound-attenuating chamber. The left pinna and cartilaginous ear canal were removed before a hollow conical speculum was positioned to give an unobstructed view of the tympanic membrane. The DPOAE probe assembly was comprised of a pair of MF1 loudspeakers (TDT) coupled to the guide tubes of an ER10B+ low noise DPOAE system (Etymotic Inc). The tip of the DPOAE probe, surrounded by a rubber gasket was positioned within the speculum. Stimuli were generated and DPOAE responses recorded using a RZ6 multifunction processor (TDT), under the control of BioSigRZ software (TDT). Continuous f1 and f2 tone stimuli were generated and presented via different MF1 drivers. Frequencies for f2 were set to match some of the ABR tone-pip frequencies used (6, 12, 18, 24 and 30 kHz). The f2 tone was presented at a frequency of 1.2 x f1, and a level 10 dB SPL lower than f1. Sound pressure levels of the f2 stimulus ranged from -10 dB to 75 dB in 5 dB steps. ER10B+ microphone signals during stimulus presentation were captured for online Fast Fourier Transformation (FFT). From each FFT trace recorded, the 2f1-f2 DPOAE amplitude, the mean noise-floor amplitude and two-times the standard deviation (SD) of the noise-floor mean were calculated. These values, plotted across stimulus level, produce plots from which the threshold of the DPOAE was defined as the lowest stimulus level where the DPOAE amplitude exceeded 2 SDs above the recording noise-floor.

#### EP recordings

The positive potential within the scala media was measured as previously described (Steel and Barkway, 1989, Ingham et al., 2016). A reference electrode (Ag-AgCl pellet) was positioned under the skin of the neck. A small hole was made in the basal turn lateral wall, and the tip of a 150 mM KCl-filled glass micropipette was inserted into the scala media. The EP was recorded as the differential potential between the tip of the glass electrode and the reference electrode, using a custom-built electrometer.

#### Preparation of cochleae and immunofluorescence microscopy

The inner ear was dissected out from the auditory bulla, a small hole was made in the cochlear apex, and the tissue was fixed in 4% paraformaldehyde (PFA) in phosphate-buffered saline (PBS; pH7.4) for 1 hour at room temperature with rotation. Following 3x 10-minute washes in PBS, the inner ear was decalcified in 4.13% EDTA (pH7.4) in PBS for 72 hours at 4 C. For postnatal mice (P1, P7 and P12) the inner ear was fixed in 4% PFA-PBS for 20-45 minutes and decalcified for 12-24 hours depending on age. All decalcified tissue was washed 3x in PBS for 10-minutes, except P1 tissue which was processed without decalcification.

For cochlear cryosections, the vestibular system was carefully removed and decalcified cochleae samples were cryoprotected with a sucrose titration then embedded in 1% low-melting point agarose in 18% sucrose-PBS solution and mounted on cryostat chucks for snap-freezing in liquid nitrogen. Mid-modiolar sections were cut at 14μm onto poly-L-lysine coated slides using a CM1860 cryostat (Leica). For organ of Corti wholemount preparations, decalcified cochleae samples were microdissected into 4 half-coils (apex; mid-apical; mid-basal and basal) using an adaptation of the Eaton-Peabody Laboratories protocol as previously described (Laboratories, 2017, Nolan et al., 2022). Cryosections were permeabilised and blocked with 10% normal goat, horse or donkey serum as appropriate in 0.2% Triton-X 100 in PBS for 1 hour at room temperature. Wholemount preparations were permeabilised with 5% Tween-20 in PBS for 1 hour at room temperature and blocked with 10% normal goat or horse serum, 0.3-0.5% Triton-X 100 in PBS for 2 hours at room temperature.

Primary antibodies were diluted in the permeabilising and blocking solution and incubated at 4 °C (cryosections) or room temperature (wholemount preparations) overnight. For adult ribbon synapse immunolabelling experiments, the primary antibodies were diluted with a 4:1 dilution of the permeabilising and blocking solution in PBS. After 3x 10-minute washes in PBS, secondary antibodies were incubated for 1-hour at room temperature in the dark. In some experiments, 10nM Phalloidin-Atto 647N (Sigma-Aldrich, Gillingham, U.K.) was added to the secondary antibody solution to label *f-*actin for visualisation of hair cell stereocilia or CellMask™ Orange (Invitrogen, #C10045) a fluorescent plasma membrane dye, diluted 1:5000 in 0.3% Triton-X 100 with PBS, was added as an extra 5-minute room temperature incubation after the secondary antibody step to visualise the myelin sheaths. Following final 3x 10-minute washes in PBS, samples were slide-mounted and coverslipped in Fluoromount-G™ (Invitrogen, #00-4958-02) or VECTASHIELD Vibrance® antifade mounting medium with DAPI (Vector Laboratories, #H-1800-2) to label cell nuclei. Imaging was performed with a Zeiss LSM 700 confocal microscope (Oberkochen, Germany) using EC Plan-Neofluar 10x/0.3 NA M27, EC Plan-Neofluar 20x/0.5 NA M27, and Plan-Apochromat 63x/1.4 NA oil M27 objectives with Zen 3.0 SR software (Black edition). For image display, equivalent linear adjustments were applied uniformly across the image where necessary to aid clarity.

#### Antibodies

##### Primary antibodies

Mouse (IgG1) anti-CtBP2 (1:400, BD Biosciences, #612044); mouse (IgG2a) anti-GluR2 (1:200, Millipore, #MAB397); rabbit (IgG) anti-MyosinVIIa (1:200, Proteus Biosciences, #25-6790); chicken (IgY) anti-Neurofilament heavy (NF-H) polypeptide (1:500, abcam, #ab4680); goat (IgG) anti-ChAT (1:200, EMD Millipore, #AB144P); mouse (IgG2a) anti-βIII-Tubulin (1:200, BioLegend, #801202); goat (IgG) anti-Peripherin (1:1000, Everest, #EB12405); mouse (IgG1ĸ) anti-Caspr (1:200, EMD Millipore, #MABN69 Clone K65/35; mouse (IgG2aĸ) Ank-G (1:200, EMD Millipore, #MABN466 N106/36).

##### Secondary antibodies

Goat anti-mouse IgG1 Alexa Fluor™ 568 (1:500; #A21124); goat anti-mouse IgG2a Alexa Fluor™ 488 (1:500; #A21131); goat anti-rabbit IgG Alexa Fluor™ 633 (1:500; #A21070); goat anti-rabbit IgG Alexa Fluor™ 405 (1:300; #A31556); goat anti-chicken IgY Alexa Fluor™ 488 (1:500; #A11039); donkey anti-mouse IgG Alexa Fluor™ 568 (1:1000; #A10037); donkey anti-rabbit IgG Alexa Fluor™ 647 (1:500; #A31573); donkey anti-goat IgG Alexa Fluor™ 488 (#1:1000; A11055) – all Invitrogen.

#### Hair cell counts, synapse analysis and quantification of myelinated fibres

Hair cells, IHC ribbon synapses and myelinated auditory fibres were quantified from organ of Corti wholemount preparations at 8kHz (mid-apical) and 24kHz (mid-basal) cochlear regions according to the mouse tonotopic frequency-place map (Muller et al., 2005). For hair cells, data was acquired from 101.6 x 76.6 µm regions of interest (ROI) generated from maximum intensity projections of confocal z-stacks collected at 0.5µm plane intervals with a Plan-Apochromat 63x/1.4 NA oil M27 objective. Hair cells were identified by Myosin7a and Phalloidin labelling and manually counted. For the IHC ribbon synapses, quantitative analysis at P60 was performed on 8-10 IHCs captured within an 84 x 25 µm ROI generated from the same maximum intensity projections used in hair cell quantification. Paired synapses were identified by manually counting the colocalised CtBP2 and GluA2 puncta within the ROI and dividing by the number of IHCs. At P12, quantitative analysis was performed on 10-12 IHCs captured within a 101.6 x 44 µm maximum intensity projection collected at 0.35µm plane intervals with a Plan-Apochromat 63x/1.4 NA oil M27 objective Counting the individual GluA2 puncta proved challenging in the cKO mice due to the reduction in the intensity of the GluA2 immunolabelling. Therefore, we focused our analysis primarily on the CtBP2 immunolabelling. First we determined the total number of CtBP2-labelled ribbons per IHC, then each ribbon was carefully examined in 2D from the maximum intensity projection using FIJI software in conjunction with the corresponding 3D projection using ZEN 3.0 SR software (Black edition) to determine whether it was an orphan or colocalised with a GluA2 puncta. All counting was performed using the Cell-Counter plugin with Fiji software (v 1.53g). For the myelinated fibres, the relative abundance was measured from a 50 x 59 µm ROI generated from maximum intensity projections of confocal z-stacks collected at 0.5µm plane intervals with an EC Plan-Neofluor 20x/0.5 NA M27 objective. Myelinated fibres were identified from the CellMask™ Orange fluorescent signal in the mid-red channel which was quantified from the ROI as the sum of pixel intensities using Fiji software (v1.53g).

#### SGNs counts

SGNs were quantified from maximum intensity projections of confocal z-stacks collected at 1µm plane intervals from midmodiolar cochlear cryosections. Images were acquired from basal, mid, and apical coils with an EC Plan-Neofluor 20x/0.5 NA M27 objective with 0.9 zoom. SGNs were identified by βIII-Tubulin labelling in conjunction with DAPI to identify the cell nuclei and manually counted within Rosenthal’s canal for each cochlear coil. To distinguish Type I SGNs from Type II, neurons that stained positive for βIII-Tubulin in the absence of Peripherin labelling were identified as Type I, whereas those that stained positive for Peripherin were identified as Type II. The cross-sectional area of Rosenthal’s canal was acquired by manually outlining the boundary of the canal from the maximum intensity projection using the freehand selection tool in Fiji (v1.53g) with the analyse > measure tool.

#### Tissue preparation and basolateral membrane recordings in IHCs

The mouse cochlea was dissected out from both male and female mice in an extracellular solution composed of (in mM): 135 NaCl, 5.8 KCl, 1.3 CaCl_2_, 0.9 MgCl_2_, 0.7 NaH_2_PO_4_, 5.6 D-glucose, 10 HEPES-NaOH. Amino acids, vitamins and sodium pyruvate (2 mM) were added from concentrates (Thermo Fisher Scientific, UK). The pH was adjusted to 7.48 with 1M NaOH (osmolality ∼308 mOsm kg^-1^). Once dissected, the cochleae were transferred to a microscope chamber and immobilised via a nylon mesh attached to a stainless-steel ring. The microscope chamber, which was continuously perfused with the above extracellular solution using a peristaltic pump (Cole-Palmer, UK), was then mounted on the stage of an upright microscopes (Olympus BX51; Nikon Eclipse FN1) with Nomarski Differential Interference Contrast (DIC) optics (60x water immersion objective) and a 15x eyepiece. All recordings were performed from apical-coil IHCs.

Patch pipettes were pulled from soda glass capillaries with a typical resistance in extracellular solution of 2-3 MΩ. Patch electrodes were coated with surf wax (Mr Zoggs SexWax, USA) to reduce the electrode capacitance. Data was acquired using an Optopatch amplifier (Cairn Research Ltd, UK), which was controlled by pClamp software using a Digidata 1440A (Molecular Devices, USA).

Potassium current recordings were performed at room temperature (20-24°C) and using the following intracellular solution (in mM): 131 KCl, 3 MgCl2, 1 EGTA-KOH, 5 Na2ATP, 5 HEPES-KOH, 10 Na-phosphocreatine (pH was adjusted with 1M KOH to 7.28; 294 mOsm kg-1). Recordings were low-pass filtered at 2.5 kHz (8-pole Bessel), sampled at 5 kHz and stored on a computer for off-line analysis (Clampfit, Molecular Devices; Origin 2021: OriginLab, USA). Membrane potentials under voltage-clamp were corrected off-line for the residual series resistance *R*_s_, which was normally compensated by 80%, and the liquid junction potential (LJP) of −4 mV, which was measured between electrode and bath solutions.

Calcium currents and real-time changes in membrane capacitance (*ΔC*_m_) were measured at body temperature (35-37°C). The intracellular solution for these experiments was (in mM): 110 Cs-glutamate, 20 CsCl2, 3 MgCl2, 1 EGTA-CsOH, 5 Na2ATP, 5 HEPES-KOH, 10 Na-phosphocreatine, 0.3 guanosine triphosphate (pH was adjusted with 1M CsOH to 7.28; 294 mOsm kg-1). In addition, the extracellular solution contained 30 mM TEA and 15 mM 4-AP to block the K^+^ currents and allow for the analysis of Ca^2+^ currents (*I_Ca_*) during the voltage step.

For *ΔC*_m_, a 4 kHz sine wave of 13 mV RMS was applied to IHCs from the membrane potential of -81 mV and was interrupted for the duration of the voltage step. The capacitance signal was filtered at 250 Hz and sampled at 5 kHz. *ΔC*_m_ was measured by averaging the *C*_m_ trace over a 200 ms period following the voltage step and subtracting the pre-pulse baseline. Calcium currents were corrected for the voltage drop across *R*_s_ and a liquid junction potential of –11 mV, measured between electrode and bath solutions.

#### SEM

The inner ear was dissected out from the auditory bulla, a small hole was made in the cochlear apex and the cochlea was gently perfused via the round window with fixative containing 2% PFA, 2.5% glutaraldehyde in 0.1M cacodylate buffer with 3mM CaCl_2_. Inner ears were then fixed for 2 hours at room temperature before being decalcified in 4% EDTA (pH7.4) in 0.1M cacodylate buffer for 48 hours at 4°C. The organ of Corti was dissected from the decalcified samples and post-fixed in 1% buffered osmium tetroxide. Samples were then processed through the osmium-thiocarbohydrazide repeated procedure (Davies and Forge, 1987) and dehydrated through an ethanol series, critical point dried, mounted and sputter coated with 5nm platinum. Images were acquired by secondary electron detection with a JEOL 6700F scanning electron microscope operating at 5kV.

#### TEM

Cochleae were fixed and decalcified as for SEM. Post-fixation, cochleae were embedded in an Epon resin (TAAB 812, TAAB, U.K.) Blocks were cut to a midmodiolar depth with an ultramicrotome. 100nm sections were taken and prepared for TEM imaging on a JEOL 1400Flash operating at 120kV. Any samples from which a classic midmodiolar orientation with intact basal, mid and apical coils could not be obtained were excluded. TEM images of the myelinated auditory nerve fibres at 12,000-15,000x magnification were acquired from the osseous spiral lamina (OSL) in the apical coil at the closest nerve bundle to the habenula perforata (HP). Axons were selected at random and the number of myelin lamellae and the thickness of the myelin sheath were quantified as shown and described in Fig. 6E.

#### RNA preparation, bulk RNA-Sequencing and Gene Ontology (GO) analysis

Auditory bullae from P1 female *Esrrg-*cKO mice and *Esrrg^fl/fl^* littermate controls were dissected in RNA*later*™ solution (Invitrogen, #AM7021) within the same 3-hour window, 2 hours after lights-on, to minimise any potential circadian effects. The vestibular system and thin bony capsule covering the cochlea were carefully removed. The soft tissue from both the left and right cochlea from one animal was pooled to make one biological replicate. Total RNA was isolated and purified using the SPLIT RNA Extraction Kit (Lexogen, #008.48) and the RNA concentration and quality were determined using a Nanodrop® spectrophotometer (Thermo Fisher Scientific, #ND-1000). Five biological replicates were prepared per genotype. RNA integrity (RIN) and quantification were assessed further using the RNA Nano 6000 Assay Kit with the Bioanalyzer 2100 system (Agilent Technologies, CA, USA), a mean RIN value of 9.6 ±0.6 was obtained from all samples. mRNA library construction and sequencing were performed by Novogene-Europe (Cambridge Science Park, U.K.) using poly A enrichment and the Illumina NovoSeq 6000 sequencer with paired-end 150bp reads, approximately, 39-49 million paired-end reads were obtained per library. Paired-end, clean reads were mapped to the mouse reference genome (mm10) using STAR (v2.5) software. HTSeq (v0.61) was used to determine the number of reads mapped to each gene. Gene expression levels were estimated from FPKM values (Reads Per Kilobase of exon model per Million mapped reads) (Mortazavi et al., 2008) and differential gene expression (DEG) analysis was performed using the DESeq2 R package (v2_1.6.3) (Anders and Huber, 2010). Gene expression thresholds were determined by an FPKM>1 and p-values were adjusted for multiple testing using Benjamini and Hochberg’s approach for controlling the False Discovery Rate. DEGs with an adjusted p-value (padj) <0.05 were considered significant. To increase stringency and facilitate the extraction of meaningful biological relevance from the data, we applied a filter of ≥ ±1.5 fold-change (≥ ±log2 0.585) to our DEGs, which is consistent with previous reports for *Esrrg* (Alaynick et al., 2007, Dufour et al., 2007). GO enrichment analysis of DEGs that met our inclusion criteria was performed with ClusterProfiler (v2.4.3). GO terms with *padj* <0.05 were considered significantly enriched. Filtering and subsequent customisation of results was performed using the cloud-based bioinformatic tool, NovoSmart (Novogene).

#### In-silico motif discovery and Ingenuity Pathway Analysis

The iRegulon (v1.3) plug-in (Janky et al., 2014) was used with Cytoscape (v3.9.1) software (Shannon et al., 2003) to predict putative binding sites for *Esrrg.* DEGs that met our cut-off criteria were submitted to iRegulon as a gene list and a scan for putative binding sites for *Esrrg* centred around 20kb of the transcription start site was run using a maximum false discovery rate of 0.001 on motif similarity with an enrichment score threshold of 2.0. Ingenuity Pathway Analysis (IPA; Qiagen) was used to identify the Canonical Signalling Pathways associated with the DEGs using the default stringent settings (Kramer et al., 2014). Significance was assessed using a right-tailed Fisher’s Exact Test, with *p*<0.05 (-log *p*=1.3).

#### qPCR analysis

RNA was prepared from P1 cochlear tissue as already described and 900ng was used in cDNA synthesis with the Omniscript® RT Kit (QIAGEN®, #205113) supplemented with random primers (Promega, #C1181), and RNaseOut™ Ribonuclease Inhibitor (Invitrogen, #10777019). Real-Time quantitative PCR was performed with TaqMan™ gene expression assays *Esrrg:* Mm00516267_m1, *Gria2:* Mm00442822_m1, *Grip2:* Mm01183453_m1, *Kcnj3:* Mm00434618_m1, *Lypd1:* Mm00513929_m1, *Myo7a:* Mm01274015_m1, *Olig1:* Mm00497537_s1, *Slc6a11*: Mm00556476_m1, *Slc17a7:* Mm00812886_m1, *Spock3:* Mm00470030_m1, *Syt7:* Mm00444502_m1, and *Tmem132d:* Mm00556406_m1 with the SsoAdvanced Universal Probes Supermix (#1725281, Bio-Rad) on a Bio-Rad CFX Connect™ Real-Time System. cDNA reactions (20ul) were diluted 1:3 and 1µl was used per qPCR reaction. Relative quantification of genes of interest was performed using the 2-ΔΔCt method with *Gapdh* as the endogenous control (#Mm99999915_g1, Applied Biosystems). 3 biological replicates were run per genotype and all samples were run as technical triplicates.

#### Statistics

In most experiments, analysis was performed on the sex-combined sample group (both males and females) before being stratified by sex to assess for potential sex-specific effects. The Shapiro-Wilk test or Kolmogorov-Smirnov (KS) test were used to assess Normality. Pairwise comparisons of mean data between genotypes were performed using a two-tailed unpaired Student’s *t*-test with the Holm-Šídák’s correction for multiple testing where applicable. A Mann-Whitney U test was used when data were not normally distributed. For multiple comparisons, a two-way analysis of variance (ANOVA) was used with Tukey’s multiple correction. All manual quantitative analysis (synapse, hair cell, myelin and SGN counts) including most of the auditory recordings were performed blind to genotype and sex. A second researcher validated the ABR threshold calls and subsets of the manual quantitative analysis. Statistical comparisons were made using GraphPad Prism (V9.1.0 and above).

## Supporting information

Supplementary Data_ALL (except Table S3)

## Data Availability

The RNA-Seq data generated in this study are available in the Gene Expression Omnibus (GEO) under accession number GSE329565.

## Author Contributions

S.V.S.: formal analysis, investigation, validation, visualisation, writing—review and editing.

N.I.: formal analysis, investigation, validation, visualisation, writing—original draft - methodology for ABR, DPOAE and EP recordings, writing—review and editing.

R.R.M.: formal analysis, investigation, visualisation, writing—review and editing.

A.J.C.: formal analysis, investigation, visualisation.

S.L.J.: formal analysis, investigation, funding acquisition, visualisation.

D.A.: formal analysis, investigation, visualisation.

A.B.: investigation, writing—review and editing

K.S.: investigation, funding acquisition, writing—review and editing.

W.M.: formal analysis, investigation, visualisation; funding acquisition, writing—original draft - methodology and results for single-hair cell electrophysiology, writing—review and editing.

K.P.S.: funding acquisition, project administration, writing—review and editing.

L.S.N.: conceptualisation, formal analysis, funding acquisition, investigation, project administration, resources, supervision, validation, visualisation, writing—original draft, writing—review and editing.

## Acknowledgements

This work was supported by a Wellcome Trust Career Development Award (225443/Z/22/Z), a King’s Prize Fellowship funded by King’s College London and the Wellcome Trust Institutional Strategic Support Fund, and a UCL Neuroscience Development Fund Award, all to L.S.N. Additional support was provided by an Action on Hearing Loss Flexi Grant (AHL-F84) to L.S.N and N.I., and the following awards: Wellcome Trust (300 350/Z/23/Z) to A.J.C.; BBSRC (BB/X000567/1) to S.L.J.; a Royal National Institute for Deaf People and Dunhill Medical Trust Fellowship (PA24) to K.E.S.; Wellcome Trust (224326/Z/21/Z) to W.M.; and Wellcome Trust (221769/Z/20/Z) to K.P.S.

The authors would like to thank the UCL Ear Institute Microscopy Unit for technical support with electron microscopy.

