## Supplementary Data_ALL (except Table S3) for "Estrogen-related receptor gamma is required for normal auditory innervation and is essential for hearing"

### Supplementary Table S1

| | | ***p(adj)*** | | | | | | | | | |
|---|---|---|---|---|---|---|---|---|---|---|---|
| | | **Stimulus / Frequency (kHz)** | | | | | | | | | |
| **Age** | **Comparison** | **Clk** | **3** | **6** | **8** | **12** | **18** | **24** | **30** | **36** | **42** |
| **2 weeks** | Pooled (Males & Females): *Esrrg-cKO* mice Vs control mice | 0.348709 | 0.348709 | **0.001238** | **0.000519** | **0.000688** | **0.024354** | **0.029164** | **0.024703** | **0.000019** | 0.348709 |
| **4 weeks** | Pooled (Males & Females): *Esrrg* -cKO mice Vs control mice | **0.000001** | **<0.000001** | **<0.000001** | **<0.000001** | **<0.000001** | **0.002401** | **0.000047** | 0.067487 | 0.770503 | 0.865725 |
| **9 weeks** | Pooled (Males & Females): *Esrrg* -cKO mice Vs control mice | **0.000094** | **<0.000001** | **<0.000001** | **<0.000001** | **0.000592** | **0.002039** | **0.002039** | **0.004381** | 0.059715 | 0.230968 |
| **2 & 4 weeks** | Pooled (Males & Females): Control mice 2 weeks Vs 4 weeks | **<0.000001** | **0.000009** | **0.005231** | **0.000298** | **0.007847** | 0.904852 | 0.964035 | 0.73825 | 0.964035 | 0.904852 |
| **4 & 9 weeks** | Pooled (Males & Females): Control mice 4 weeks Vs 9 weeks | 0.616648 | 0.899305 | 0.899305 | 0.899305 | 0.956765 | 0.974169 | 0.955495 | 0.974169 | 0.963995 | 0.974169 |
| **2 & 4 weeks** | Pooled (Males & Females): *Esrrg* -cKO mice 2 weeks Vs 4 weeks | **0.005767** | 0.073811 | 0.073811 | **0.017391** | **0.000055** | **0.002709** | **0.018095** | 0.073811 | **0.015221** | 0.123573 |
| **4 & 9 weeks** | Pooled (Males & Females): *Esrrg* -cKO mice 4 weeks Vs 9 weeks | 0.952787 | 0.910201 | 0.934111 | 0.934111 | 0.952787 | 0.952787 | 0.910201 | 0.495617 | 0.910201 | 0.910201 |

**Table S1.** Statistical comparisons for **Fig. 2A-C.** Mean ABR thresholds to click and pure tone stimuli in *Esrrg*$^{+/+}$ and *Esrrg*-cKO mice at different timepoints. Results of unpaired, two-tailed, *t*-test with Holm-Šídák's correction for multiple testing with significant results shown in bold.

#### Supplementary Figure S1

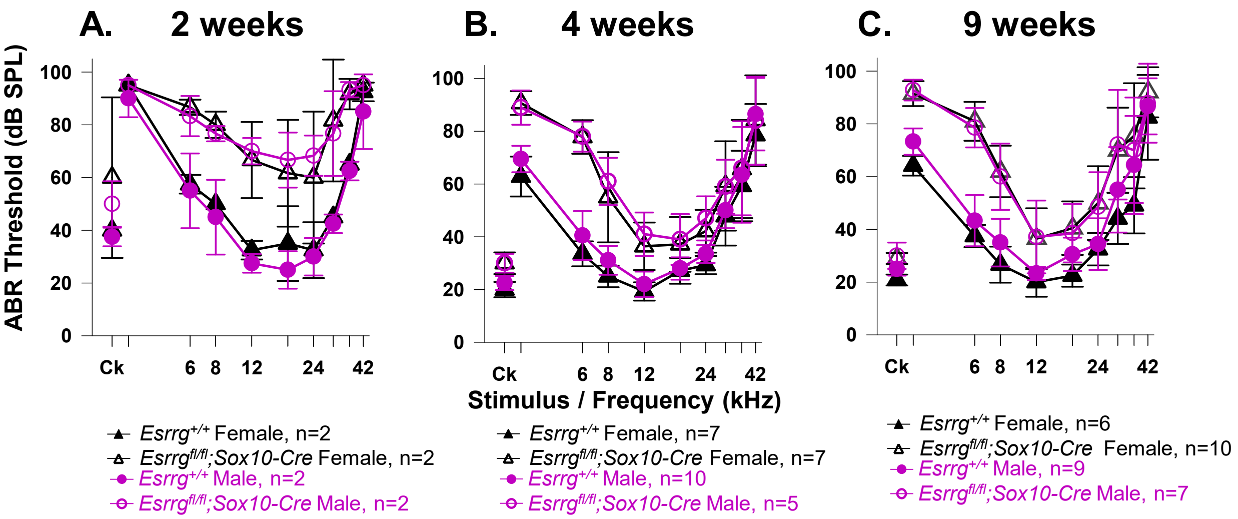

***Figure S1. Sex does not impact auditory brainstem response (ABR) thresholds in Esrrg-cKO mice.*** Mean ABR thresholds (±SD) to click and pure tone stimuli in *Esrrg*$^{+/+}$ and *Esrrg*$^{fl/fl}$*;Sox10-Cre* mice at 2 (P14), 4 (P31-P37) and 9 (P60-P64) weeks of age. Auditory thresholds in control mice did not significantly differ between the sexes at any age except 9 weeks - slightly higher thresholds were observed in males for click stimuli and at 3kHz. Auditory thresholds in *Esrrg*$^{fl/fl}$*;Sox10-Cre* mice did not significantly differ between males and females at any age. The number and sex of the mice is shown below the thresholds. Statistical comparisons: unpaired *t*-test with Holm-Šídák's correction for multiple testing (*padj:* **Table S2**).

#### Supplementary Table S2

| | | *p(adj)* | | | | | | | | | |
|---|---|---|---|---|---|---|---|---|---|---|---|
| | | Stimulus / Frequency (kHz) | | | | | | | | | |
| **Age** | **Comparison** | **Clk** | **3** | **6** | **8** | **12** | **18** | **24** | **30** | **36** | **42** |
| **2 weeks** | Males Vs Females in control mice | 0.992866 | 0.992866 | 0.992866 | 0.992866 | 0.968947 | 0.992866 | 0.992866 | 0.992866 | 0.992866 | 0.992866 |
| | Males Vs Females in *Esrrg* -cKO mice | 0.996502 | >0.999999 | 0.993946 | 0.976553 | 0.996502 | 0.996502 | 0.996502 | 0.996502 | 0.996502 | >0.999999 |
| | Males: *Esrrg* -cKO mice Vs control mice | 0.406805 | 0.470264 | 0.253256 | 0.149081 | **0.017944** | 0.111918 | 0.085211 | 0.253256 | **0.016768** | 0.470264 |
| | Females: *Esrrg* -cKO mice Vs control mice | 0.611934 | >0.999999 | **0.017748** | **0.031587** | 0.274402 | 0.611934 | 0.611934 | 0.480993 | 0.057663 | 0.611934 |
| **4 weeks** | Males Vs Females in control mice | 0.459618 | 0.304175 | 0.459618 | 0.219684 | 0.717777 | 0.940461 | 0.215511 | 0.940461 | 0.940461 | 0.717777 |
| | Males Vs Females in *Esrrg* -cKO mice | >0.999999 | 0.998374 | >0.999999 | 0.995608 | 0.988403 | 0.998374 | 0.983210 | >0.999999 | >0.999999 | >0.999999 |
| | Males: *Esrrg* -cKO mice Vs control mice | **0.002737** | **0.000165** | **0.000017** | **0.000041** | **0.000504** | **0.030685** | **0.002751** | 0.167110 | 0.943162 | 0.943162 |
| | Females: *Esrrg* -cKO mice Vs control mice | **0.001526** | **0.000021** | **<0.000001** | **0.004204** | **0.003565** | 0.317698 | **0.011178** | 0.410515 | 0.721454 | 0.721454 |
| **9 weeks** | Males Vs Females in control mice | **0.042666** | **0.021498** | 0.519988 | 0.430736 | 0.505438 | 0.131030 | 0.799756 | 0.519988 | 0.447078 | 0.799756 |
| | Males Vs Females in *Esrrg* -cKO mice | 0.998898 | 0.998898 | 0.998898 | 0.998898 | 0.998898 | 0.998898 | 0.998898 | 0.998898 | 0.998898 | 0.998898 |
| | Males: *Esrrg* -cKO mice Vs control mice | 0.114051 | **0.000007** | **0.000013** | **0.003043** | 0.068963 | 0.297136 | 0.128378 | 0.297136 | 0.819483 | 0.854698 |
| | Females: *Esrrg* -cKO mice Vs control mice | **0.000018** | **<0.000001** | **<0.000001** | **0.000016** | **0.023030** | **0.006404** | **0.030430** | **0.016744** | **0.030430** | 0.155160 |

**Table S2.** Statistical comparisons for **Fig. S1.** Mean ABR thresholds to click and pure tone stimuli in *Esrrg*-cKO mice stratified by sex. Results of unpaired, two-tailed, *t*-test with Holm-Šídák's correction for multiple testing with significant results shown in bold.

### Supplementary Figure S2

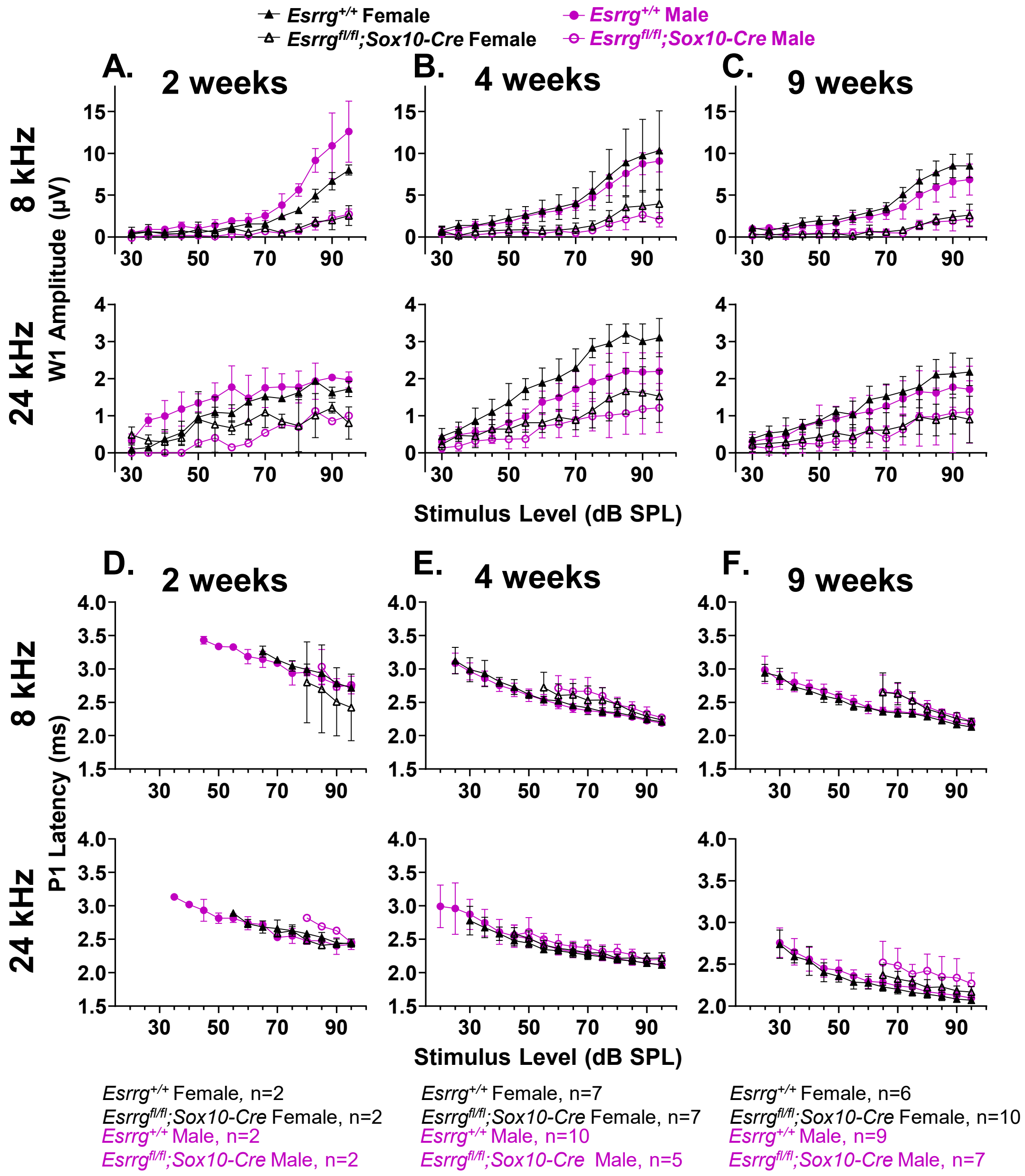

***Figure S2. The impact of sex on wave 1 amplitude and latency in Esrrg-cKO mice.*** **(A-C)** Mean wave 1 amplitude (±SD) at 8kHz and 24kHz plotted as a function of dB sound pressure level in *Esrrg*$^{+/+}$ and *Esrrg*$^{fl/fl}$*;Sox10-Cre* mice at 2, 4 and 9 weeks of age. For clarity data is shown from 30dB SPL. **(D-F)** Mean wave 1 latency (±SD) at 8kHz and 24kHz plotted as a function of dB sound pressure level in *Esrrg*$^{+/+}$ and *Esrrg*$^{fl/fl}$*;Sox10-Cre* mice at 2, 4 and 9 weeks of age. Latency is shown from the mean threshold (dB SPL) for each genotype and frequency. Data from **(A-F)** were obtained from the same mice as in **Fig. S1**.

### Supplementary Figure S3

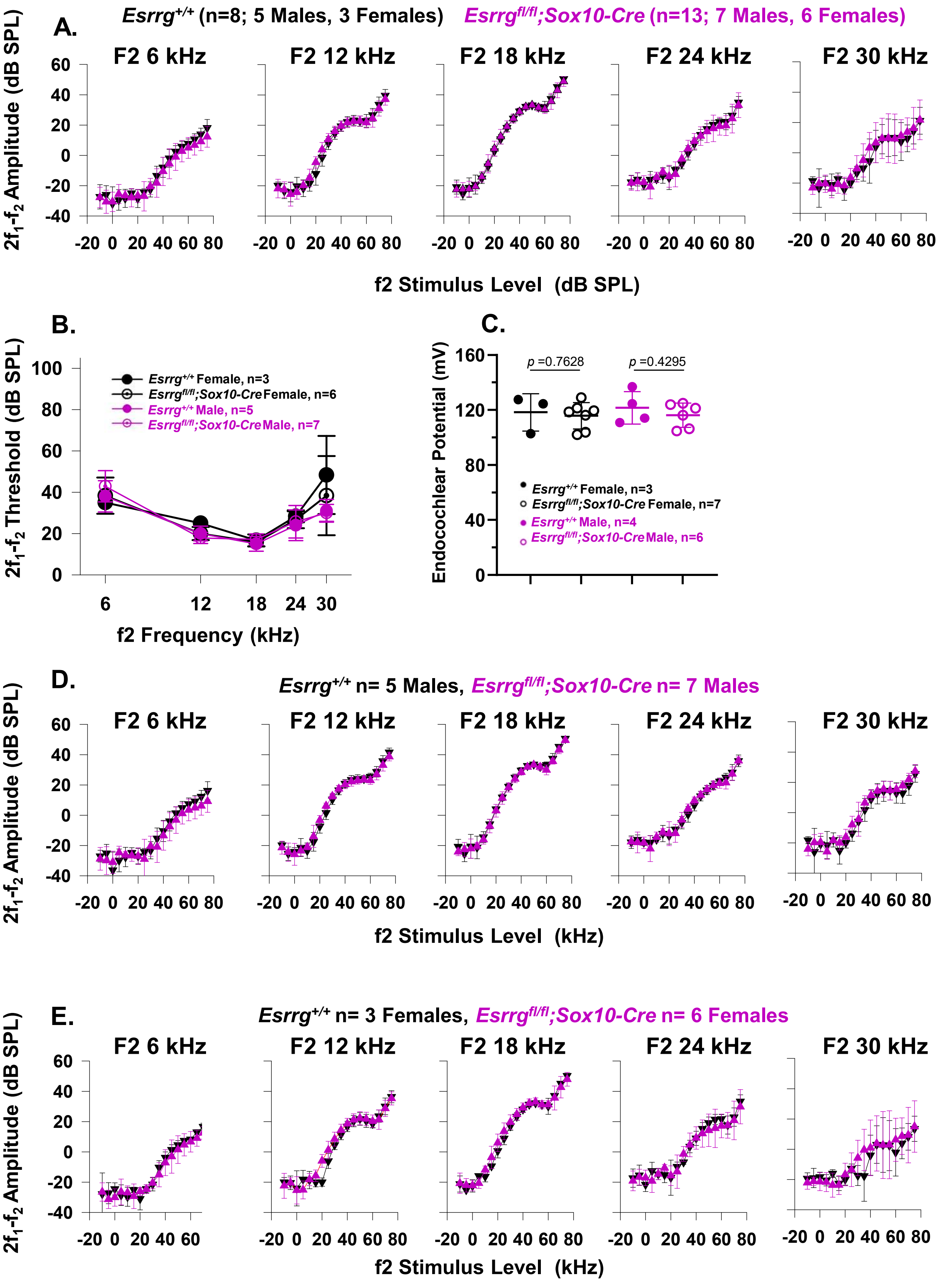

***Figure S3. Sex does not impact DPOAEs and the EP in young adult Esrrg-cKO mice.*** Mean (±SD) $2f_1$-$f_2$ DPOAE amplitudes at f2 frequencies 6kHz-30kHz plotted as a function of f2 stimulus level (dB SPL) in *Esrrg*$^{+/+}$ and *Esrrg*$^{fl/fl}$*;Sox10-Cre* mice at 9 weeks of age: **(A)** Pooled data – males and females; **(D)** males only; **(E)** females only.

**(B)** Sex-stratified analysis to show mean (±SD) $2f_1$-$f_2$ DPOAE thresholds plotted as a function of f2 frequency (kHz) in *Esrrg*$^{+/+}$ and *Esrrg*$^{fl/fl}$*;Sox10-Cre* mice at 9 weeks of age (P57-P62). Statistical comparisons: 2-way ANOVA with Tukey's multiple correction; Males: $p$ = 0.4460; Females: $p$ = 0.3375.

**(C)** Sex-stratified analysis to show EP recordings from *Esrrg*$^{+/+}$ and *Esrrg*$^{fl/fl}$*;Sox10-Cre* mice at 9 weeks of age. Data is plotted as the mean (±SD). Statistical comparisons: unpaired *t*-test. Recordings were taken from the same mice used in (B); number and sex are shown below the thresholds.

#### Supplementary Figure S4

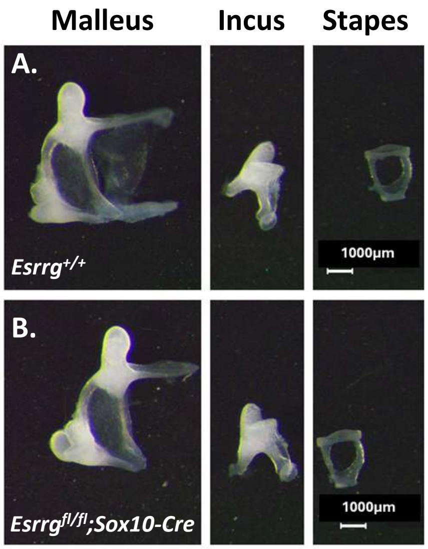

***Figure S4: Normal morphology of the auditory ossicles in Esrrg-cKO mice.*** Morphological characteristics of the auditory ossicles in **(A)** *Esrrg*$^{+/+}$ and **(B)** *Esrrg*$^{fl/fl}$*;Sox10-Cre* mice at P29 to show normal appearance of the malleus, incus and stapes. Representative images from 2-3 mice per genotype per sex are shown.

#### Supplementary Figure S5

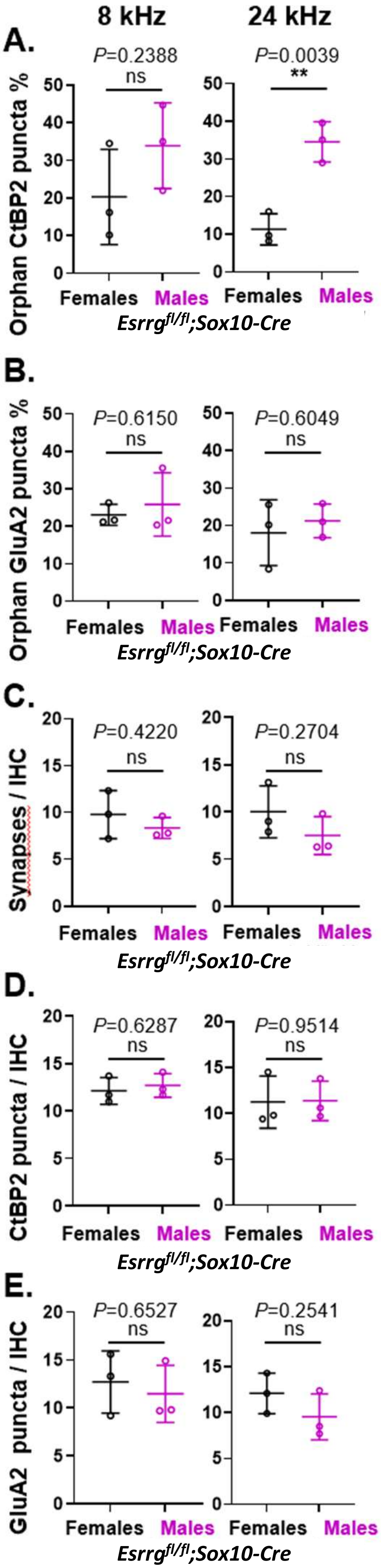

***Figure S5. The impact of sex on the IHC ribbon synapse in adult Esrrg-cKO mice. Expanded analysis from Figure 3.*** (A) Orphan ribbons and (B) orphan GluA2 densities in female and male *Esrrg*$^{fl/fl}$*;Sox10-Cre* mice expressed as a percentage of the total number of ribbons / GluA2 densities, respectively. (C) Number of paired synapses, (D) CtBP2 puncta and (E) GluA2 puncta per IHC in female and male *Esrrg*$^{fl/fl}$*;Sox10-Cre* mice. Data is sampled from 8-10 IHCs in 3 *Esrrg*$^{fl/fl}$*;Sox10-Cre* mice per sex. All data is plotted as mean values ± SD; **p<0.01 unpaired *t*-test.

#### Supplementary Figure S6

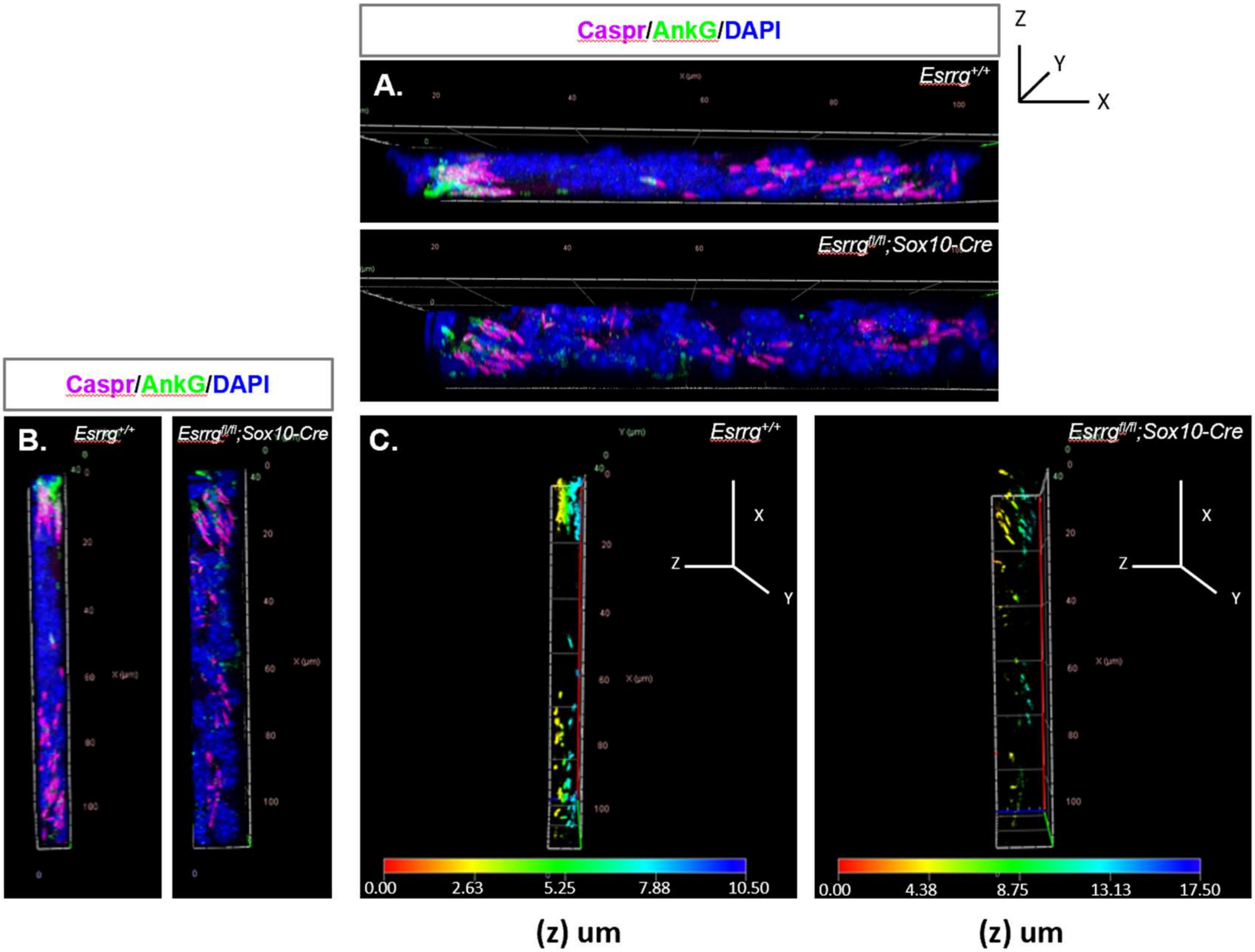

***Figure S6: 3D-renderings of first heminode configuration in Esrrg-cKO mice.***

(A-C) 3D-renderings of confocal z-stacks of apical coil cochlear cryosections from adult *Esrrg*$^{+/+}$ and *Esrrg*$^{fl/fl}$*;Sox10-Cre* mice immunolabelled with antibodies to the heminodal proteins - Caspr (Magenta) and AnkG (Green). Different configurations are shown as indicated by the x,y,z plane. (C) shows the same image as (B) with depth coding where the channels are colour coded based on the z-axis.

#### Supplementary Figure S7

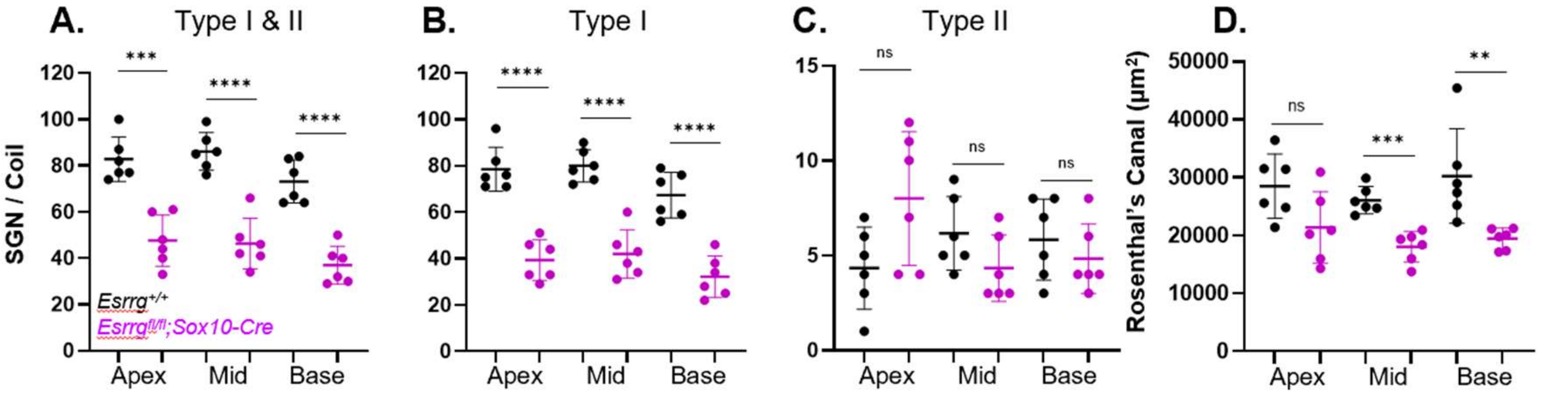

***Figure S7: Adult Esrrg-cKO mice have fewer Type I SGNs encased in smaller Rosenthal's canals.*** (A-D) Quantification of SGNs at P29 in *Esrrg$^{fl/fl}$;Sox10-Cre* mice compared to controls across basal, mid and apical cochlear coils shows reduced numbers of Type I SGNs in *Esrrg$^{fl/fl}$;Sox10-Cre* mice (B) in conjunction with a smaller Rosenthal's canal (D). The Type II counts (C) did not differ by number. Data was acquired from 3 mice per genotype per sex and is plotted as mean values ± SD; *$p$<0.05; **$p$<0.01; ***$p$<0.001; ****$p$<0.0001 unpaired t-test**.**

#### Supplementary Figure S8

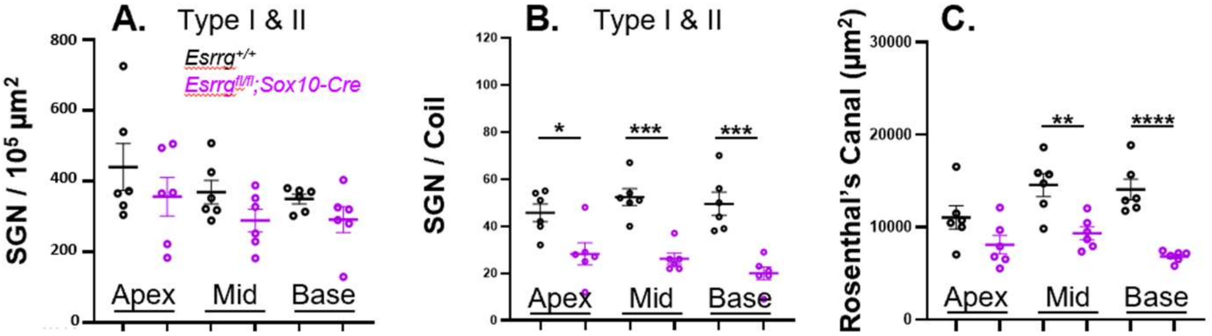

***Figure S8. Loss of SGNs in Esrrg-cKO mice from early development.*** (A-C) Quantification of Type I and II SGNs at P1 in *Esrrg$^{fl/fl}$;Sox10-Cre* mice compared to controls across basal, mid and apical cochlear coils shows reduced numbers of SGNs in *Esrrg$^{fl/fl}$;Sox10-Cre* mice (B) in conjunction with a smaller Rosenthal's canal (C). The density of the SGNs (A) shows a trend towards reduction in *Esrrg$^{fl/fl}$;Sox10-Cre* mice, but this did not reach significance. Data was acquired from 3 mice per genotype per sex and is plotted as mean values ± SD; *$p$<0.05; **$p$<0.01; ***$p$<0.001; ****$p$<0.0001 unpaired t-test**.**

#### Supplementary Figure S9

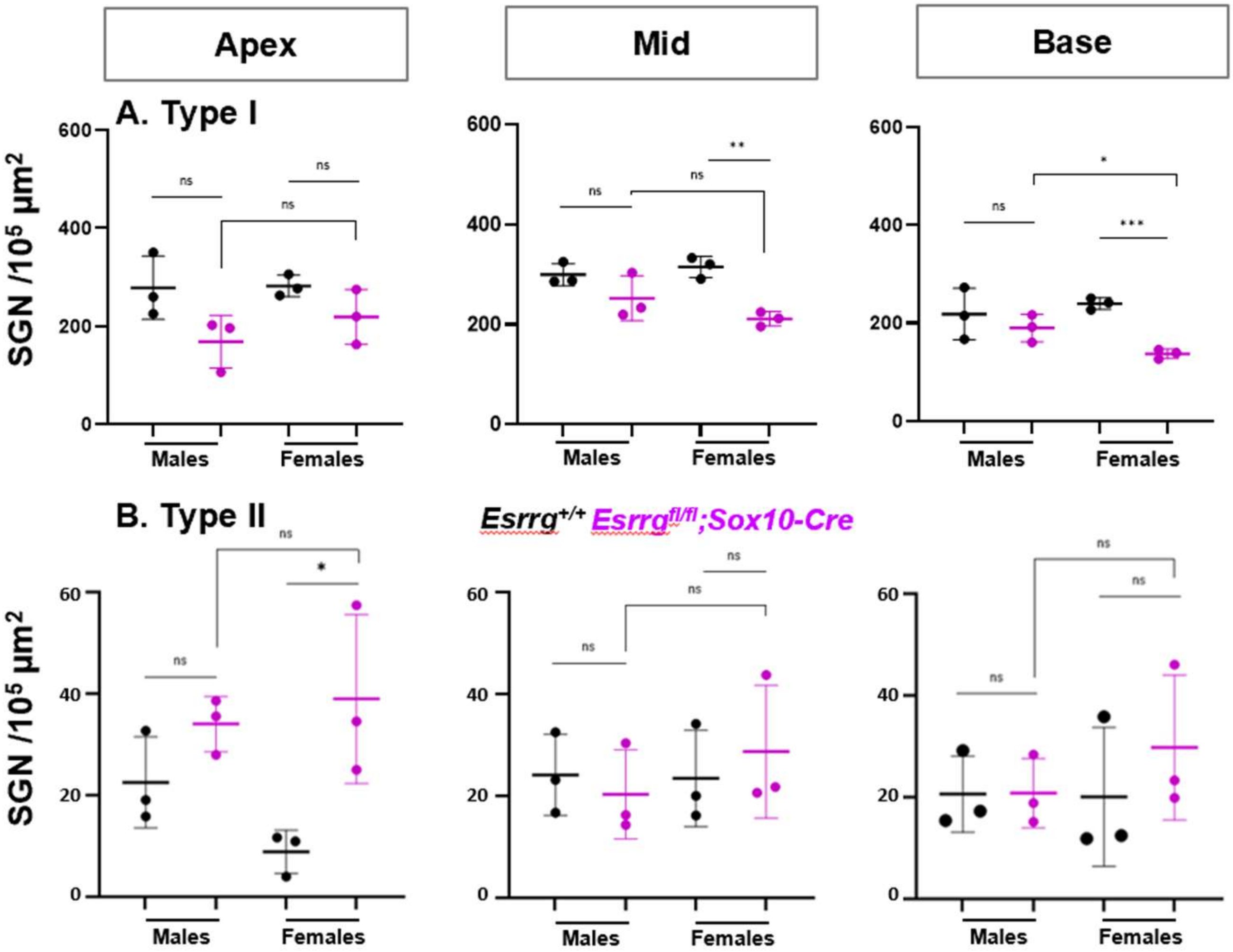

***Figure S9: The impact of sex on SGN density in Esrrg-cKO mice.*** (**A**-**B**) SGN density at P29 in Type **I** (**A**) and Type **II** (**B**) SGNs in *Esrrg*$^{fl/fl}$*;Sox10-Cre* mice compared to controls. Data is shown for apical, mid and basal cochlear coils, stratified by sex and acquired from 3 mice per genotype per sex, plotted as mean values ± SD; *p<0.05; **p<0.01; ***p<0.001 unpaired t-test.

#### Supplementary Figure S10

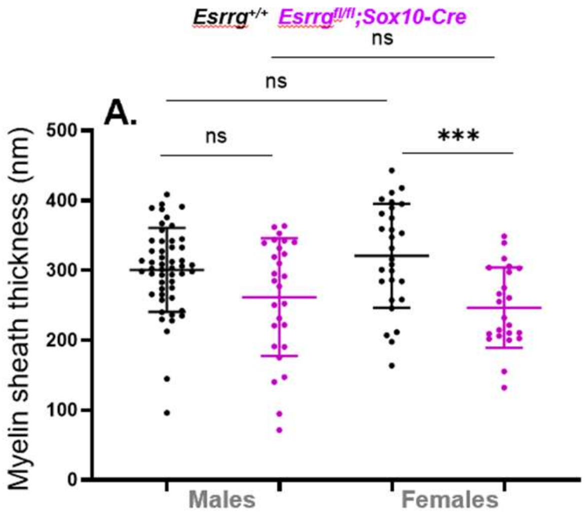

***Figure S10: Sex-stratified analysis of the myelin ultrastructure in Esrrg-cKO mice.*** Myelin sheath thickness (A) and number of myelin lamellae (B) in *Esrrg*$^{+/+}$ (n=9; 4 females, 5 males) versus *Esrrg*$^{fl/fl}$*;Sox10-Cre* (n=7; 4 females, 3 males) mice. Metrics were obtained from at least 8 apical auditory nerve fibres per sample. Statistical analysis was conducted using an unpaired t-test *(***$p$<0.01; ****$p$<0.0001)

### Supplementary Table S3

***Supplementary Table S3: DEGs in Esrrg*-cKO *mice.*** Only DEGs that met our inclusion criteria (log2FoldChange ± ≥ 0.585; padj<0.05) are listed.

**ABOVE TABLE IS AN EXCEL FILE**

#### Supplementary Table S4

| Gene | log2FoldChange | padj |
|---|---|---|
| *Abl1* | 0.181 | 0.599372 |
| *Actb* | -0.152 | 0.719921 |
| *B2m* | -0.029 | 0.963340 |
| *Casc3* | 0.061 | 0.883059 |
| *Cdkn1a* | -0.205 | 0.446983 |
| *Cdkn1b* | -0.112 | 0.673399 |
| *Eif2b1* | -0.090 | 0.810487 |
| *Elf1* | 0.196 | 0.399874 |
| *Gadd45a* | 0.003 | 0.996400 |
| *Gapdh* | -0.103 | 0.790998 |
| *Gusb* | 0.001 | 0.998499 |
| *Hmbs* | -0.127 | 0.806837 |
| *Hmbs* | -0.127 | 0.806837 |
| *Hprt* | -0.340 | 0.347877 |
| *Ipo8* | 0.181 | 0.634339 |
| *Mrpl19* | -0.151 | 0.681699 |
| *mt-Atp6* | -0.063 | 0.914880 |
| *Pes1* | -0.021 | 0.949466 |
| *Pgk1* | -0.253 | 0.418374 |
| *Polr2a* | 0.297 | 0.552559 |
| *Pop4* | -0.059 | 0.903040 |
| *Ppia* | -0.191 | 0.644420 |
| *Psmc4* | -0.137 | 0.686474 |
| *Pum1* | 0.178 | 0.487417 |
| *Rpl30* | -0.131 | 0.801315 |
| *Rpl37a* | -0.280 | 0.623957 |
| *Rplp0* | 0.043 | 0.944810 |
| *Rplp2* | -0.177 | 0.766828 |
| *Rps17* | -0.178 | 0.722166 |
| *Tbp* | -0.142 | 0.700138 |
| ***Tfrc*** | **-0.722** | **2.430000E-11** |
| *Tubb3* | -0.386 | 0.309131 |
| *Ubc* | -0.316 | 0.789598 |
| *Ywhaz* | -0.016 | 0.965822 |

***Supplementary Table S4: Comparison of gene expression profiles for common housekeeping genes in Esrrg-cKO mice.*** Housekeeping genes were selected from the TaqMan Array Mouse Endogenous Control Panel (#4426701; Applied Biosystems). Housekeeping genes which were significantly dysregulated in *Esrrg*$^{fl/fl}$*;Sox10-Cre* mice are shown in bold (log2FoldChange ± ≥ 0.585; padj<0.05). Results for the pan-neuronal marker *Tubb3* which encodes β-III Tubulin are also shown. Genes are ordered alphabetically.

#### Supplementary Figure S11

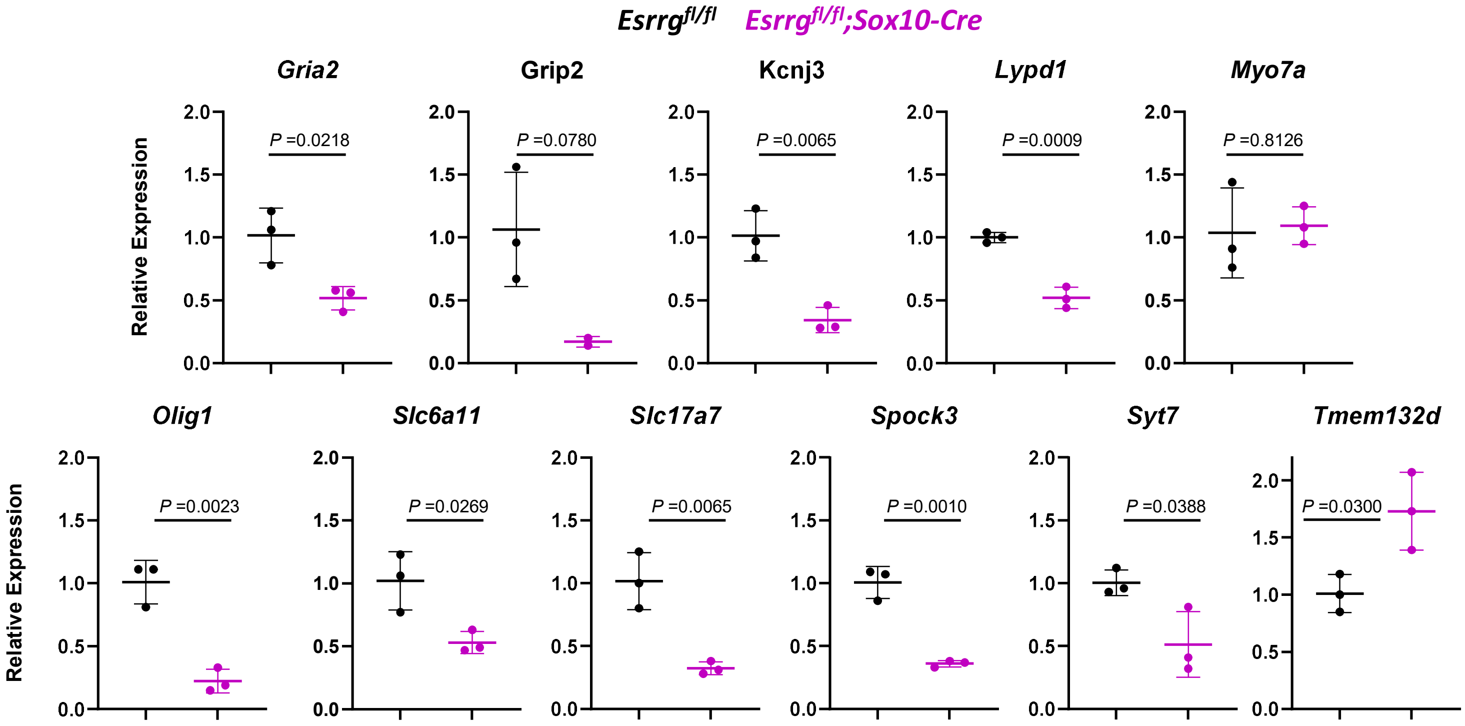

***Figure S11: qPCR analysis of putative Esrrg targets in the early postnatal cochlea. Expanded analysis from Figure 7.*** qPCR analysis was conducted on a subset of putative *Esrrg targets* using RNA isolated from P1 cochleae from *Esrrg*$^{fl/fl}$*;Sox10-Cre* mice and sex-matched *Esrrg*$^{fl/fl}$ littermate controls (female). Relative quantification levels were adjusted to *Gapdh* as an endogenous control and normalized to *Esrrg*$^{fl/fl}$ levels. Data are presented as mean (±SD); statistical comparisons: unpaired *t*-test with a Holm-Sidak correction for multiple testing. The *Myo7a* gene, a marker for the sensory hair cells, was also included as a negative control as this gene was not found to vary in our RNA-Seq analysis (Fig.7A,B), nor was this gene predicted to contain binding sites for *Esrrg* from our iRegulon analysis (Fig.7C). *Gria2,* which encodes the GluA2 protein, although not a hit from our iRegulon bioinformatic analysis (Fig.7C) was also included due to the sparse labelling of the IHC ribbon post-synaptic density with an anti-GluA2 antibody at P12 (Fig.3H-K).

#### Supplementary Table S5

| Marker | log2foldchange | padj | Gene name | Ref |
|---|---|---|---|---|
| Oligodendrocyte precursor cells / regulator of peripheral glial development. | 0.146 | 0.672276 | *Sox10* | Kuhlbrodt, Herbarth et al. 1998, Watanabe, Takeda et al. 2000, Britsch, Goerich et al. 2001. |
| Early marker of cochlear satellite and Schwann cell glia. | -0.068 | 0.917159 | *Sox2* | Smith, Murphy et al. 2021. |

**Supplementary Table S5:** ***Early markers of cochlear peripheral glia development are not dysregulated in Esrrg-cKO mice.*** Results of bulk RNA-Seq analysis from whole cochleae tissue from female *Esrrg*$^{fl/fl}$ (n=5) and *Esrrg*$^{fl/fl}$*;Sox10-Cre* (n=5) mice at P1. Data is shown for two common cochlear glia markers.

#### Supplementary Table S6

| Gene | Direction | Putative Target |
|---|---|---|
| ***Aqp4*** | **DOWN** | **Yes** |
| *Clrn1* | DOWN | No |
| *Dio3* | UP | No |
| ***Dlx1*** | **DOWN** | **Yes** |
| ***Hcn2*** | **DOWN** | **Yes** |
| *Lrrtm3* | DOWN | No |
| ***Mafb*** | **DOWN** | **Yes** |
| *Otos* | DOWN | No |
| *Prkcb* | DOWN | No |
| *Rimbp2* | DOWN | No |
| ***Scn11a*** | **UP** | **Yes** |
| *Sema3e* | DOWN | No |

***Supplementary Table S6: DEGs in Esrrg-cKO mice implicated in hearing.*** DEGs that met our inclusion criteria (log2FoldChange ± ≥ 0.585; padj<0.05) were compared to a manually curated list of genes implicated in hearing from mouse and/or human studies (https://doi.org/10.5281/zenodo.15705587) and cross-referenced against the DEGs that contained putative *Esrrg*-binding sites (Fig.7C).
