## Supplementary Data_Table S3 for "Estrogen-related receptor gamma is required for normal auditory innervation and is essential for hearing"

| gene_id | log2FoldChange | pvalue | padj | gene_name | gene_description |
|---|---|---|---|---|---|
| ENSMUSG00000054162 | -1.413696809 | 4.28E-28 | 8.05E-24 | Spock3 | sparc/osteonectin, cwcv and kazal-like domains proteoglycan 3 [Source:MGI Symbol;Acc:MGI:1920152] |
| ENSMUSG00000046160 | -2.749646709 | 2.55E-27 | 2.40E-23 | Olig1 | oligodendrocyte transcription factor 1 [Source:MGI Symbol;Acc:MGI:1355334] |
| ENSMUSG00000032503 | -1.52727208 | 2.04E-25 | 1.28E-21 | Arpp21 | cyclic AMP-regulated phosphoprotein, 21 [Source:MGI Symbol;Acc:MGI:107562] |
| ENSMUSG00000017400 | -1.350529512 | 8.39E-23 | 3.95E-19 | Stac2 | SH3 and cysteine rich domain 2 [Source:MGI Symbol;Acc:MGI:2144518] |
| ENSMUSG00000033047 | 0.806524014 | 9.89E-22 | 3.72E-18 | Eif3l | eukaryotic translation initiation factor 3, subunit L [Source:MGI Symbol;Acc:MGI:2386251] |
| ENSMUSG00000015134 | -1.162734081 | 5.62E-21 | 1.76E-17 | Aldh1a3 | aldehyde dehydrogenase family 1, subfamily A3 [Source:MGI Symbol;Acc:MGI:1861722] |
| ENSMUSG00000040653 | -1.220892932 | 7.54E-20 | 2.03E-16 | Ppp1r14c | protein phosphatase 1, regulatory inhibitor subunit 14C [Source:MGI Symbol;Acc:MGI:1923392] |
| ENSMUSG00000027834 | -0.913962064 | 3.42E-18 | 8.05E-15 | Serpini1 | serine (or cysteine) peptidase inhibitor, clade I, member 1 [Source:MGI Symbol;Acc:MGI:1194506] |
| ENSMUSG00000024743 | -1.105044273 | 3.56E-17 | 7.44E-14 | Syt7 | synaptotagmin VII [Source:MGI Symbol;Acc:MGI:1859545] |
| ENSMUSG00000003657 | -1.667487911 | 4.03E-17 | 7.59E-14 | Calb2 | calbindin 2 [Source:MGI Symbol;Acc:MGI:101914] |
| ENSMUSG00000056306 | -0.829968507 | 3.71E-16 | 6.34E-13 | Sertm1 | serine rich and transmembrane domain containing 1 [Source:MGI Symbol;Acc:MGI:3607715] |
| ENSMUSG00000000632 | -1.141254665 | 4.54E-16 | 7.12E-13 | Sez6 | seizure related gene 6 [Source:MGI Symbol;Acc:MGI:104745] |
| ENSMUSG00000029304 | -1.27579375 | 2.81E-15 | 4.07E-12 | Spp1 | secreted phosphoprotein 1 [Source:MGI Symbol;Acc:MGI:98389] |
| ENSMUSG00000022952 | -0.83359425 | 1.06E-14 | 1.24E-11 | Runx1 | runt related transcription factor 1 [Source:MGI Symbol;Acc:MGI:99852] |
| ENSMUSG00000032908 | -0.912564506 | 9.91E-15 | 1.24E-11 | Sgpp2 | sphingosine-1-phosphate phosphotase 2 [Source:MGI Symbol;Acc:MGI:3589109] |
| ENSMUSG00000030270 | -1.608223538 | 9.25E-15 | 1.24E-11 | Cpne9 | copine family member IX [Source:MGI Symbol;Acc:MGI:2443052] |
| ENSMUSG00000070570 | -1.122111959 | 1.27E-14 | 1.40E-11 | Slc17a7 | solute carrier family 17 (sodium-dependent inorganic phosphate cotransporter), member 7 [Source:MGI Symbol;Acc:MGI:1920211] |
| ENSMUSG00000022797 | -0.721971559 | 2.32E-14 | 2.43E-11 | Tfrc | transferrin receptor [Source:MGI Symbol;Acc:MGI:98822] |
| ENSMUSG00000008658 | -1.04269281 | 7.99E-14 | 7.91E-11 | Rbfox1 | RNA binding protein, fox-1 homolog (C. elegans) 1 [Source:MGI Symbol;Acc:MGI:1926224] |
| ENSMUSG00000097452 | 2.242507752 | 1.11E-13 | 1.04E-10 | Gm2824 | predicted gene 2824 [Source:MGI Symbol;Acc:MGI:3780995] |
| ENSMUSG00000028546 | -0.731996852 | 1.39E-13 | 1.24E-10 | Elavl4 | ELAV like RNA binding protein 4 [Source:MGI Symbol;Acc:MGI:107427] |
| ENSMUSG00000034310 | 1.052874722 | 4.81E-13 | 4.11E-10 | Tmem132d | transmembrane protein 132D [Source:MGI Symbol;Acc:MGI:3044963] |
| ENSMUSG00000038173 | -1.759373671 | 6.45E-13 | 5.28E-10 | Enpp6 | ectonucleotide pyrophosphatase/phosphodiesterase 6 [Source:MGI Symbol;Acc:MGI:2445171] |
| ENSMUSG00000052889 | -0.635389595 | 7.25E-13 | 5.69E-10 | Prkcb | protein kinase C, beta [Source:MGI Symbol;Acc:MGI:97596] |
| ENSMUSG00000044364 | -1.024755857 | 9.18E-13 | 6.91E-10 | Tmem74b | transmembrane protein 74B [Source:MGI Symbol;Acc:MGI:1918629] |
| ENSMUSG00000024411 | -1.611572518 | 1.22E-12 | 8.86E-10 | Aqp4 | aquaporin 4 [Source:MGI Symbol;Acc:MGI:107387] |
| ENSMUSG00000033039 | 1.09586388 | 1.73E-12 | 1.21E-09 | Micall1 | microtubule associated monooxygenase, calponin and LIM domain containing -like 1 [Source:MGI Symbol;Acc:MGI:105870] |
| ENSMUSG00000075707 | 3.356202471 | 3.74E-12 | 2.51E-09 | Dio3 | deiodinase, iodothyronine type III [Source:MGI Symbol;Acc:MGI:1306782] |
| ENSMUSG00000043557 | -0.980176223 | 3.99E-12 | 2.59E-09 | Mdga1 | MAM domain containing glycosylphosphatidylinositol anchor 1 [Source:MGI Symbol;Acc:MGI:1922012] |
| ENSMUSG00000042846 | -1.065235444 | 1.01E-11 | 6.34E-09 | Lrrtm3 | leucine rich repeat transmembrane neuronal 3 [Source:MGI Symbol;Acc:MGI:2389177] |
| ENSMUSG00000019772 | 4.427093305 | 1.45E-11 | 8.80E-09 | Vip | vasoactive intestinal polypeptide [Source:MGI Symbol;Acc:MGI:98933] |
| ENSMUSG00000031841 | 0.828557378 | 1.83E-11 | 1.08E-08 | Cdh13 | cadherin 13 [Source:MGI Symbol;Acc:MGI:99551] |
| ENSMUSG00000024798 | -1.273615023 | 2.50E-11 | 1.42E-08 | Htr7 | 5-hydroxytryptamine (serotonin) receptor 7 [Source:MGI Symbol;Acc:MGI:99841] |
| ENSMUSG00000033020 | 0.846010816 | 2.95E-11 | 1.63E-08 | Polr2f | polymerase (RNA) II (DNA directed) polypeptide F [Source:MGI Symbol;Acc:MGI:1349393] |
| ENSMUSG00000091722 | -1.090653247 | 3.24E-11 | 1.74E-08 | Siah3 | siah E3 ubiquitin protein ligase family member 3 [Source:MGI Symbol;Acc:MGI:2685758] |
| ENSMUSG00000058420 | -0.914644421 | 4.04E-11 | 2.11E-08 | Syt17 | synaptotagmin XVII [Source:MGI Symbol;Acc:MGI:104966] |
| ENSMUSG00000052229 | -2.28767557 | 6.42E-11 | 3.27E-08 | Gpr17 | G protein-coupled receptor 17 [Source:MGI Symbol;Acc:MGI:3584514] |
| ENSMUSG00000097545 | -1.505236898 | 7.42E-11 | 3.67E-08 | Mir124a-1hg | Mir124-1 host gene (non-protein coding) [Source:MGI Symbol;Acc:MGI:2442197] |
| ENSMUSG00000039830 | -2.040663291 | 8.22E-11 | 3.97E-08 | Olig2 | oligodendrocyte transcription factor 2 [Source:MGI Symbol;Acc:MGI:1355331] |
| ENSMUSG00000039457 | -0.799547282 | 2.10E-10 | 9.87E-08 | Ppl | periplakin [Source:MGI Symbol;Acc:MGI:1194898] |
| ENSMUSG00000045509 | -1.146528394 | 2.28E-10 | 1.05E-07 | Gpr150 | G protein-coupled receptor 150 [Source:MGI Symbol;Acc:MGI:2441872] |
| ENSMUSG00000048385 | -0.828913525 | 2.72E-10 | 1.19E-07 | Scrt1 | scratch family zinc finger 1 [Source:MGI Symbol;Acc:MGI:2176606] |
| ENSMUSG00000070695 | -1.159961746 | 2.67E-10 | 1.19E-07 | Cntnap5a | contactin associated protein-like 5A [Source:MGI Symbol;Acc:MGI:3643623] |
| ENSMUSG00000035551 | -0.936009815 | 3.75E-10 | 1.60E-07 | Igfbpl1 | insulin-like growth factor binding protein-like 1 [Source:MGI Symbol;Acc:MGI:1933198] |
| ENSMUSG00000094002 | -0.774715195 | 4.64E-10 | 1.94E-07 | Gm9866 | predicted gene 9866 [Source:MGI Symbol;Acc:MGI:3643118] |
| ENSMUSG00000035561 | -0.844740119 | 1.02E-09 | 4.16E-07 | Aldh1b1 | aldehyde dehydrogenase 1 family, member B1 [Source:MGI Symbol;Acc:MGI:1919785] |
| ENSMUSG00000029757 | -0.590324388 | 1.94E-09 | 7.77E-07 | Dync1i1 | dynein cytoplasmic 1 intermediate chain 1 [Source:MGI Symbol;Acc:MGI:107743] |
| ENSMUSG00000074622 | -0.745568856 | 2.35E-09 | 9.23E-07 | Mafb | v-maf musculoaponeurotic fibrosarcoma oncogene family, protein B (avian) [Source:MGI Symbol;Acc:MGI:104555] |
| ENSMUSG00000043850 | -0.664449165 | 4.50E-09 | 1.73E-06 | Clrn1 | clarin 1 [Source:MGI Symbol;Acc:MGI:2388124] |
| ENSMUSG00000046607 | 1.127822824 | 5.99E-09 | 2.25E-06 | Hrk | harakiri, BCL2 interacting protein (contains only BH3 domain) [Source:MGI Symbol;Acc:MGI:1201608] |
| ENSMUSG00000067879 | -1.818457657 | 6.60E-09 | 2.43E-06 | Vxn | vexin [Source:MGI Symbol;Acc:MGI:1924232] |
| ENSMUSG00000045034 | -1.182710267 | 7.13E-09 | 2.58E-06 | Ankrd34b | ankyrin repeat domain 34B [Source:MGI Symbol;Acc:MGI:2443245] |
| ENSMUSG00000033061 | -0.970331399 | 1.94E-08 | 6.90E-06 | Resp18 | regulated endocrine-specific protein 18 [Source:MGI Symbol;Acc:MGI:1098222] |
| ENSMUSG00000063531 | -0.79363805 | 3.09E-08 | 1.08E-05 | Sema3e | sema domain, immunoglobulin domain (Ig), short basic domain, secreted, (semaphorin) 3E [Source:MGI Symbol;Acc:MGI:1340034] |
| ENSMUSG00000024403 | -0.735614124 | 4.32E-08 | 1.48E-05 | Atp6v1g2 | ATPase, H+ transporting, lysosomal V1 subunit G2 [Source:MGI Symbol;Acc:MGI:1913487] |
| ENSMUSG00000054459 | -0.934852827 | 4.60E-08 | 1.55E-05 | Vsnl1 | visinin-like 1 [Source:MGI Symbol;Acc:MGI:1349453] |

| | | | | | |
|---|---|---|---|---|---|
| ENSMUSG00000056755 | -0.898558951 | 5.21E-08 | 1.72E-05 | Grm7 | glutamate receptor, metabotropic 7 [Source:MGI Symbol;Acc:MGI:1351344] |
| ENSMUSG00000027577 | -0.728741647 | 8.49E-08 | 2.71E-05 | Chrna4 | cholinergic receptor, nicotinic, alpha polypeptide 4 [Source:MGI Symbol;Acc:MGI:87888] |
| ENSMUSG00000035805 | -2.363280079 | 1.03E-07 | 3.25E-05 | Mlc1 | megalencephalic leukoencephalopathy with subcortical cysts 1 homolog (human) [Source:MGI Symbol;Acc:MGI:2157910] |
| ENSMUSG00000021852 | -1.146754878 | 1.11E-07 | 3.43E-05 | Slc35f4 | solute carrier family 35, member F4 [Source:MGI Symbol;Acc:MGI:1922538] |
| ENSMUSG00000044835 | -0.762321448 | 1.42E-07 | 4.32E-05 | Ankrd45 | ankyrin repeat domain 45 [Source:MGI Symbol;Acc:MGI:1921094] |
| ENSMUSG00000059361 | -0.940092629 | 1.54E-07 | 4.60E-05 | Nrsn2 | neurensin 2 [Source:MGI Symbol;Acc:MGI:2684969] |
| ENSMUSG00000034115 | 1.230422061 | 1.62E-07 | 4.77E-05 | Scn11a | sodium channel, voltage-gated, type XI, alpha [Source:MGI Symbol;Acc:MGI:1345149] |
| ENSMUSG00000024347 | -0.65936814 | 1.75E-07 | 5.06E-05 | Psd2 | pleckstrin and Sec7 domain containing 2 [Source:MGI Symbol;Acc:MGI:1921252] |
| ENSMUSG00000027173 | -0.775278039 | 1.88E-07 | 5.37E-05 | Depdc7 | DEP domain containing 7 [Source:MGI Symbol;Acc:MGI:2139258] |
| ENSMUSG00000033316 | -0.858217529 | 1.99E-07 | 5.52E-05 | Galnt9 | polypeptide N-acetylgalactosaminyltransferase 9 [Source:MGI Symbol;Acc:MGI:2677965] |
| ENSMUSG00000015829 | -1.47873045 | 1.97E-07 | 5.52E-05 | Tnr | tenascin R [Source:MGI Symbol;Acc:MGI:99516] |
| ENSMUSG00000037977 | -0.650162156 | 2.11E-07 | 5.77E-05 | 6430571L13Rik | RIKEN cDNA 6430571L13 gene [Source:MGI Symbol;Acc:MGI:2445137] |
| ENSMUSG00000031543 | -0.861118079 | 2.65E-07 | 7.12E-05 | Ank1 | ankyrin 1, erythroid [Source:MGI Symbol;Acc:MGI:88024] |
| ENSMUSG00000021803 | -0.741700521 | 3.03E-07 | 7.91E-05 | Cdhr1 | cadherin-related family member 1 [Source:MGI Symbol;Acc:MGI:2157782] |
| ENSMUSG00000028773 | -1.040166538 | 3.01E-07 | 7.91E-05 | Fabp3 | fatty acid binding protein 3, muscle and heart [Source:MGI Symbol;Acc:MGI:95476] |
| ENSMUSG00000026824 | -1.458838035 | 4.12E-07 | 0.000104882 | Kcnj3 | potassium inwardly-rectifying channel, subfamily J, member 3 [Source:MGI Symbol;Acc:MGI:104742] |
| ENSMUSG00000028217 | 0.792470056 | 4.60E-07 | 0.00011522 | Cdh17 | cadherin 17 [Source:MGI Symbol;Acc:MGI:1095414] |
| ENSMUSG00000021196 | -0.675819754 | 4.65E-07 | 0.00011522 | Pfkp | phosphofructokinase, platelet [Source:MGI Symbol;Acc:MGI:1891833] |
| ENSMUSG00000021194 | -0.902897865 | 4.95E-07 | 0.000120881 | Chga | chromogranin A [Source:MGI Symbol;Acc:MGI:88394] |
| ENSMUSG00000032355 | -0.950690964 | 5.41E-07 | 0.00013047 | Mlip | muscular LMNA-interacting protein [Source:MGI Symbol;Acc:MGI:1916892] |
| ENSMUSG00000027350 | -0.749566077 | 6.82E-07 | 0.000158464 | Chgb | chromogranin B [Source:MGI Symbol;Acc:MGI:88395] |
| ENSMUSG00000034336 | -0.668779401 | 6.97E-07 | 0.000159907 | Ina | internexin neuronal intermediate filament protein, alpha [Source:MGI Symbol;Acc:MGI:96568] |
| ENSMUSG00000022044 | -0.748186248 | 7.63E-07 | 0.000171534 | Stmn4 | stathmin-like 4 [Source:MGI Symbol;Acc:MGI:1931224] |
| ENSMUSG00000097023 | -1.48014767 | 7.66E-07 | 0.000171534 | Mir9-3hg | Mir9-3 host gene [Source:MGI Symbol;Acc:MGI:2142071] |
| ENSMUSG00000027674 | -0.697153069 | 7.83E-07 | 0.000173332 | Pex5l | peroxisomal biogenesis factor 5-like [Source:MGI Symbol;Acc:MGI:1916672] |
| ENSMUSG00000026163 | -0.867176505 | 9.16E-07 | 0.000200488 | Sphkap | SPHK1 interactor, AKAP domain containing [Source:MGI Symbol;Acc:MGI:1924879] |
| ENSMUSG00000026610 | 0.623631587 | 1.23E-06 | 0.000260487 | Esrrg | estrogen-related receptor gamma [Source:MGI Symbol;Acc:MGI:1347056] |
| ENSMUSG00000075224 | -1.08970122 | 1.86E-06 | 0.000385408 | Lrrc55 | leucine rich repeat containing 55 [Source:MGI Symbol;Acc:MGI:2685197] |
| ENSMUSG00000042115 | -1.078644081 | 1.90E-06 | 0.000389552 | Klhdc8a | kelch domain containing 8A [Source:MGI Symbol;Acc:MGI:2442630] |
| ENSMUSG00000021587 | 1.174734074 | 2.25E-06 | 0.000450059 | Pcsk1 | proprotein convertase subtilisin/kexin type 1 [Source:MGI Symbol;Acc:MGI:97511] |
| ENSMUSG00000031654 | -0.882988553 | 2.37E-06 | 0.000468547 | Cbln1 | cerebellin 1 precursor protein [Source:MGI Symbol;Acc:MGI:88281] |
| ENSMUSG00000079045 | -1.545552921 | 2.43E-06 | 0.000477194 | Prox1os | prospero homeobox 1, opposite strand [Source:MGI Symbol;Acc:MGI:4937200] |
| ENSMUSG00000032523 | -0.617170579 | 2.61E-06 | 0.000507267 | Hhatl | hedgehog acyltransferase-like [Source:MGI Symbol;Acc:MGI:1922020] |
| ENSMUSG00000025370 | -1.017029155 | 2.72E-06 | 0.000522489 | Cdh9 | cadherin 9 [Source:MGI Symbol;Acc:MGI:107433] |
| ENSMUSG00000060257 | -0.763036098 | 2.84E-06 | 0.000539724 | Scrt2 | scratch family zinc finger 2 [Source:MGI Symbol;Acc:MGI:2139287] |
| ENSMUSG00000002190 | -0.617302807 | 3.05E-06 | 0.000567413 | Clgn | calmegin [Source:MGI Symbol;Acc:MGI:107472] |
| ENSMUSG00000022435 | -1.908572255 | 3.36E-06 | 0.000620835 | Upk3a | uroplakin 3A [Source:MGI Symbol;Acc:MGI:98914] |
| ENSMUSG00000040797 | -0.791647225 | 3.93E-06 | 0.000717685 | Iqsec3 | IQ motif and Sec7 domain 3 [Source:MGI Symbol;Acc:MGI:2677208] |
| ENSMUSG00000030307 | -0.851561619 | 4.24E-06 | 0.000752227 | Slc6a11 | solute carrier family 6 (neurotransmitter transporter, GABA), member 11 [Source:MGI Symbol;Acc:MGI:95630] |
| ENSMUSG00000018589 | -0.932925561 | 4.33E-06 | 0.000761071 | Glra2 | glycine receptor, alpha 2 subunit [Source:MGI Symbol;Acc:MGI:95748] |
| ENSMUSG00000016349 | -0.796913337 | 4.95E-06 | 0.000847285 | Eef1a2 | eukaryotic translation elongation factor 1 alpha 2 [Source:MGI Symbol;Acc:MGI:1096317] |
| ENSMUSG00000023000 | -0.642014857 | 6.49E-06 | 0.001099497 | Dhh | desert hedgehog [Source:MGI Symbol;Acc:MGI:94891] |
| ENSMUSG00000061762 | 1.931255609 | 7.82E-06 | 0.001313453 | Tac1 | tachykinin 1 [Source:MGI Symbol;Acc:MGI:98474] |
| ENSMUSG00000047324 | -0.862487988 | 8.90E-06 | 0.001469171 | 4931429P17Rik | RIKEN cDNA 4931429P17 gene [Source:MGI Symbol;Acc:MGI:1918221] |
| ENSMUSG00000031688 | -0.69564753 | 9.21E-06 | 0.001506446 | Pou4f2 | POU domain, class 4, transcription factor 2 [Source:MGI Symbol;Acc:MGI:102524] |
| ENSMUSG00000097156 | -1.704219876 | 9.39E-06 | 0.001522443 | Gm3764 | predicted gene 3764 [Source:MGI Symbol;Acc:MGI:3781938] |
| ENSMUSG00000050840 | -0.951001458 | 9.94E-06 | 0.001585958 | Cdh20 | cadherin 20 [Source:MGI Symbol;Acc:MGI:1346069] |
| ENSMUSG00000030844 | -0.683860457 | 1.18E-05 | 0.001807168 | Rgs10 | regulator of G-protein signalling 10 [Source:MGI Symbol;Acc:MGI:1915115] |
| ENSMUSG00000115529 | -1.332813817 | 1.18E-05 | 0.001807168 | 9630013A20Rik | RIKEN cDNA 9630013A20 gene [Source:MGI Symbol;Acc:MGI:2442953] |
| ENSMUSG00000053963 | -0.632404369 | 1.25E-05 | 0.001902378 | Stum | mechanosensory transduction mediator [Source:MGI Symbol;Acc:MGI:2138735] |
| ENSMUSG00000029420 | -0.73939361 | 1.29E-05 | 0.001936721 | Rimbp2 | RIMS binding protein 2 [Source:MGI Symbol;Acc:MGI:2443235] |
| ENSMUSG00000114289 | -0.775676209 | 1.76E-05 | 0.002600002 | Gm48453 | predicted gene, 48453 [Source:MGI Symbol;Acc:MGI:6097966] |
| ENSMUSG00000027360 | 1.04285295 | 2.04E-05 | 0.002976989 | Hdc | histidine decarboxylase [Source:MGI Symbol;Acc:MGI:96062] |
| ENSMUSG00000053166 | -0.724286255 | 2.14E-05 | 0.00307001 | Cdh22 | cadherin 22 [Source:MGI Symbol;Acc:MGI:1341843] |
| ENSMUSG00000110631 | 0.745234299 | 2.22E-05 | 0.003161957 | Gm42047 | predicted gene, 42047 [Source:MGI Symbol;Acc:MGI:5624932] |
| ENSMUSG00000060240 | -0.818504539 | 2.23E-05 | 0.003161957 | Cend1 | cell cycle exit and neuronal differentiation 1 [Source:MGI Symbol;Acc:MGI:1929898] |
| ENSMUSG00000108601 | -0.589388947 | 2.56E-05 | 0.003548253 | Gm44645 | predicted gene 44645 [Source:MGI Symbol;Acc:MGI:5753221] |

| | | | | | |
|---|---|---|---|---|---|
| ENSMUSG00000007944 | -0.771894333 | 2.56E-05 | 0.003548253 | Ttc9b | tetratricopeptide repeat domain 9B [Source:MGI Symbol;Acc:MGI:1920282] |
| ENSMUSG00000026344 | -0.663387108 | 2.79E-05 | 0.003835186 | Lypd1 | Ly6/Plaur domain containing 1 [Source:MGI Symbol;Acc:MGI:1919835] |
| ENSMUSG00000047344 | -0.859712312 | 3.15E-05 | 0.004294114 | Lancl3 | LanC lantibiotic synthetase component C-like 3 (bacterial) [Source:MGI Symbol;Acc:MGI:2443335] |
| ENSMUSG00000027071 | -0.62401614 | 3.24E-05 | 0.004392716 | P2rx3 | purinergic receptor P2X, ligand-gated ion channel, 3 [Source:MGI Symbol;Acc:MGI:1097160] |
| ENSMUSG00000041911 | -0.658068408 | 3.55E-05 | 0.004777107 | Dlx1 | distal-less homeobox 1 [Source:MGI Symbol;Acc:MGI:94901] |
| ENSMUSG00000104157 | -1.719263055 | 3.94E-05 | 0.00517906 | Gm38101 | predicted gene, 38101 [Source:MGI Symbol;Acc:MGI:5611329] |
| ENSMUSG00000037843 | -0.72806751 | 4.24E-05 | 0.005507086 | Vstm2l | V-set and transmembrane domain containing 2-like [Source:MGI Symbol;Acc:MGI:2685537] |
| ENSMUSG00000020774 | -0.723678002 | 4.64E-05 | 0.005942968 | Aspa | aspartoacylase [Source:MGI Symbol;Acc:MGI:87914] |
| ENSMUSG00000105954 | -2.336649524 | 4.76E-05 | 0.006055997 | Gm42793 | predicted gene 42793 [Source:MGI Symbol;Acc:MGI:5662930] |
| ENSMUSG00000005716 | -1.203570427 | 4.98E-05 | 0.006290287 | Pvalb | parvalbumin [Source:MGI Symbol;Acc:MGI:97821] |
| ENSMUSG00000005045 | -0.607220386 | 5.02E-05 | 0.006296357 | Chd5 | chromodomain helicase DNA binding protein 5 [Source:MGI Symbol;Acc:MGI:3036258] |
| ENSMUSG00000036480 | -2.617082177 | 5.12E-05 | 0.00637853 | Prss56 | protease, serine 56 [Source:MGI Symbol;Acc:MGI:1916703] |
| ENSMUSG00000026424 | -0.892221364 | 5.30E-05 | 0.006566767 | Gpr37l1 | G protein-coupled receptor 37-like 1 [Source:MGI Symbol;Acc:MGI:1928503] |
| ENSMUSG00000033029 | 0.856932476 | 5.34E-05 | 0.006570121 | 1700088E04Rik | RIKEN cDNA 1700088E04 gene [Source:MGI Symbol;Acc:MGI:1920774] |
| ENSMUSG00000026312 | -0.978026383 | 5.46E-05 | 0.006672734 | Cdh7 | cadherin 7, type 2 [Source:MGI Symbol;Acc:MGI:2442792] |
| ENSMUSG00000086096 | -0.734069287 | 6.10E-05 | 0.0073063 | Gm12688 | predicted gene 12688 [Source:MGI Symbol;Acc:MGI:3650434] |
| ENSMUSG00000068696 | -0.593738411 | 6.42E-05 | 0.007603485 | Gpr88 | G-protein coupled receptor 88 [Source:MGI Symbol;Acc:MGI:1927653] |
| ENSMUSG00000033981 | -0.806815715 | 7.72E-05 | 0.008915026 | Gria2 | glutamate receptor, ionotropic, AMPA2 (alpha 2) [Source:MGI Symbol;Acc:MGI:95809] |
| ENSMUSG00000044349 | -1.210612054 | 7.90E-05 | 0.009054939 | Snhg11 | small nucleolar RNA host gene 11 [Source:MGI Symbol;Acc:MGI:2441845] |
| ENSMUSG00000011154 | -0.966834033 | 9.08E-05 | 0.010104271 | Cfap161 | cilia and flagella associated protein 161 [Source:MGI Symbol;Acc:MGI:1922806] |
| ENSMUSG00000035849 | -0.682589857 | 0.000103818 | 0.011164295 | Krt222 | keratin 222 [Source:MGI Symbol;Acc:MGI:2442728] |
| ENSMUSG00000113737 | 2.759267973 | 0.000142317 | 0.014797073 | BB123696 | expressed sequence BB123696 [Source:MGI Symbol;Acc:MGI:2145475] |
| ENSMUSG00000038242 | 0.844238447 | 0.000147426 | 0.015243983 | Aox4 | aldehyde oxidase 4 [Source:MGI Symbol;Acc:MGI:1919122] |
| ENSMUSG00000019874 | -0.678228534 | 0.000153966 | 0.015833232 | Fabp7 | fatty acid binding protein 7, brain [Source:MGI Symbol;Acc:MGI:101916] |
| ENSMUSG00000096918 | -1.416559874 | 0.000163367 | 0.016618354 | Gm16863 | predicted gene, 16863 [Source:MGI Symbol;Acc:MGI:4439787] |
| ENSMUSG00000073988 | -0.795374952 | 0.000167694 | 0.016966838 | Ttpa | tocopherol (alpha) transfer protein [Source:MGI Symbol;Acc:MGI:1354168] |
| ENSMUSG00000015467 | -0.67721351 | 0.000199885 | 0.019360489 | Egfl8 | EGF-like domain 8 [Source:MGI Symbol;Acc:MGI:1932094] |
| ENSMUSG00000027168 | -1.55037655 | 0.000199834 | 0.019360489 | Pax6 | paired box 6 [Source:MGI Symbol;Acc:MGI:97490] |
| ENSMUSG00000020672 | -0.753743761 | 0.000206082 | 0.019686577 | Sntg2 | syntrophin, gamma 2 [Source:MGI Symbol;Acc:MGI:1919541] |
| ENSMUSG00000033854 | -0.785974173 | 0.000229331 | 0.02168729 | Kcnk10 | potassium channel, subfamily K, member 10 [Source:MGI Symbol;Acc:MGI:1919508] |
| ENSMUSG00000044519 | -1.85172363 | 0.000240054 | 0.022475514 | Zfp488 | zinc finger protein 488 [Source:MGI Symbol;Acc:MGI:2686052] |
| ENSMUSG00000020331 | -0.627332474 | 0.000254767 | 0.023618023 | Hcn2 | hyperpolarization-activated, cyclic nucleotide-gated K+ 2 [Source:MGI Symbol;Acc:MGI:1298210] |
| ENSMUSG00000026069 | 0.898328342 | 0.000262468 | 0.02412628 | Il1rl1 | interleukin 1 receptor-like 1 [Source:MGI Symbol;Acc:MGI:98427] |
| ENSMUSG00000058740 | -0.836490468 | 0.000295491 | 0.026107263 | Kcnt1 | potassium channel, subfamily T, member 1 [Source:MGI Symbol;Acc:MGI:1924627] |
| ENSMUSG00000046159 | -0.715074837 | 0.000310971 | 0.026844786 | Chrm3 | cholinergic receptor, muscarinic 3, cardiac [Source:MGI Symbol;Acc:MGI:88398] |
| ENSMUSG00000047897 | -1.315813468 | 0.000310038 | 0.026844786 | Ripply2 | ripply transcriptional repressor 2 [Source:MGI Symbol;Acc:MGI:2685968] |
| ENSMUSG00000097518 | -0.659603105 | 0.000319888 | 0.027488492 | Gm26694 | predicted gene, 26694 [Source:MGI Symbol;Acc:MGI:5477188] |
| ENSMUSG00000051515 | -0.679770427 | 0.000326205 | 0.027822374 | Fam181b | family with sequence similarity 181, member B [Source:MGI Symbol;Acc:MGI:1930951] |
| ENSMUSG00000085562 | 1.450021468 | 0.000342278 | 0.028672417 | 2610028E06Rik | RIKEN cDNA 2610028E06 gene [Source:MGI Symbol;Acc:MGI:1919645] |
| ENSMUSG00000030098 | -1.067843276 | 0.000348789 | 0.028672417 | Grip2 | glutamate receptor interacting protein 2 [Source:MGI Symbol;Acc:MGI:2681173] |
| ENSMUSG00000092448 | -1.324488669 | 0.00034046 | 0.028672417 | Gm20387 | predicted gene 20387 [Source:MGI Symbol;Acc:MGI:5141852] |
| ENSMUSG00000027434 | -1.940172916 | 0.000347294 | 0.028672417 | Nkx2-2 | NK2 homeobox 2 [Source:MGI Symbol;Acc:MGI:97347] |
| ENSMUSG00000041468 | -2.274163592 | 0.000347539 | 0.028672417 | Gpr12 | G-protein coupled receptor 12 [Source:MGI Symbol;Acc:MGI:101909] |
| ENSMUSG00000027581 | -0.687609814 | 0.000357516 | 0.029125972 | Stmn3 | stathmin-like 3 [Source:MGI Symbol;Acc:MGI:1277137] |
| ENSMUSG00000019235 | -0.614856542 | 0.000366299 | 0.029458868 | Rps6kl1 | ribosomal protein S6 kinase-like 1 [Source:MGI Symbol;Acc:MGI:2443413] |
| ENSMUSG00000040972 | -0.599130625 | 0.000388928 | 0.030685027 | Igsf21 | immunoglobulin superfamily, member 21 [Source:MGI Symbol;Acc:MGI:2681842] |
| ENSMUSG00000032224 | -0.622486057 | 0.000418219 | 0.031993732 | Fam81a | family with sequence similarity 81, member A [Source:MGI Symbol;Acc:MGI:1924136] |
| ENSMUSG00000029875 | -0.683089554 | 0.00044637 | 0.033600979 | Ccdc184 | coiled-coil domain containing 184 [Source:MGI Symbol;Acc:MGI:2146066] |
| ENSMUSG00000116504 | -0.983008141 | 0.00044611 | 0.033600979 | I730030J21Rik | RIKEN cDNA I730030J21 gene [Source:MGI Symbol;Acc:MGI:3588279] |
| ENSMUSG00000025425 | -0.639189062 | 0.000459478 | 0.034311604 | St8sia5 | ST8 alpha-N-acetyl-neuraminide alpha-2,8-sialyltransferase 5 [Source:MGI Symbol;Acc:MGI:109243] |
| ENSMUSG00000037224 | -0.614992378 | 0.000468501 | 0.03458305 | Zfyve28 | zinc finger, FYVE domain containing 28 [Source:MGI Symbol;Acc:MGI:2684992] |
| ENSMUSG00000055775 | 1.348622688 | 0.000475501 | 0.034818907 | Myh8 | myosin, heavy polypeptide 8, skeletal muscle, perinatal [Source:MGI Symbol;Acc:MGI:1339712] |
| ENSMUSG00000048978 | -0.675148319 | 0.000495991 | 0.036038832 | Nrsn1 | neurensin 1 [Source:MGI Symbol;Acc:MGI:894662] |
| ENSMUSG00000022468 | -0.667671128 | 0.000533436 | 0.038315766 | Endou | endonuclease, polyU-specific [Source:MGI Symbol;Acc:MGI:97746] |
| ENSMUSG00000044055 | -0.784029012 | 0.000537716 | 0.038476326 | Otos | otospiralin [Source:MGI Symbol;Acc:MGI:2672814] |
| ENSMUSG00000041923 | -0.704943557 | 0.000546064 | 0.038925689 | Nol4 | nucleolar protein 4 [Source:MGI Symbol;Acc:MGI:2441684] |
| ENSMUSG00000020887 | -1.053919273 | 0.000550602 | 0.039101069 | A230052G05Ri | RIKEN cDNA A230052G05 gene [Source:MGI Symbol;Acc:MGI:3045239] |

| | | | | | |
|---|---|---|---|---|---|
| ENSMUSG00000019785 | -0.823116383 | 0.000557865 | 0.039467897 | Clvs2 | clavesin 2 [Source:MGI Symbol;Acc:MGI:2443223] |
| ENSMUSG00000021908 | -3.519107395 | 0.000573688 | 0.039986025 | Gm6768 | predicted gene 6768 [Source:MGI Symbol;Acc:MGI:3648259] |
| ENSMUSG00000044092 | -1.262909339 | 0.000625788 | 0.042515192 | C130050O18Ri | RIKEN cDNA C130050O18 gene [Source:MGI Symbol;Acc:MGI:2442694] |
| ENSMUSG00000024935 | -0.875416267 | 0.000634043 | 0.042921065 | Slc1a1 | solute carrier family 1 (neuronal/epithelial high affinity glutamate transporter, system Xag), member 1 [Source:MGI Symbol;Acc:MGI:105083] |
| ENSMUSG00000018865 | -0.62173362 | 0.000660583 | 0.044240232 | Sult4a1 | sulfotransferase family 4A, member 1 [Source:MGI Symbol;Acc:MGI:1888971] |
| ENSMUSG00000078640 | -0.705304323 | 0.00067906 | 0.045156265 | Gm11627 | predicted gene 11627 [Source:MGI Symbol;Acc:MGI:3650659] |
| ENSMUSG00000046854 | -0.838762764 | 0.000709028 | 0.046983082 | Pip5kl1 | phosphatidylinositol-4-phosphate 5-kinase-like 1 [Source:MGI Symbol;Acc:MGI:2448520] |
| ENSMUSG00000086905 | -1.257382296 | 0.000724325 | 0.0476611 | Gm13716 | predicted gene 13716 [Source:MGI Symbol;Acc:MGI:3650432] |
| ENSMUSG00000025020 | -0.850947227 | 0.000753382 | 0.04922878 | Slit1 | slit guidance ligand 1 [Source:MGI Symbol;Acc:MGI:1315203] |
